# Immune-responsive gene-1: The mitochondrial Key to Th17 Cell Pathogenicity in CNS Autoimmunity

**DOI:** 10.1101/2023.12.24.573264

**Authors:** Mohammad Nematullah, Mena Fatma, Guoli Zhou, Faraz Rashid, Kameshwar Ayasolla, Mohammad Ejaz Ahmed, Ruicong She, Sajad Mir, Insha Zahoor, Nasrul Hoda, Ramandeep Rattan, Shailendra Giri

## Abstract

Pathogenic Th17 cells play crucial roles in CNS autoimmune diseases such as multiple sclerosis (MS), but their regulation by endogenous mechanisms remains unknown. Through RNA-seq analysis of primary brain glial cells, we identified immuno-responsive gene 1 (*Irg1*) as one of the highly upregulated gene under inflammatory conditions. Validation in the spinal cord of animals with experimental autoimmune encephalomyelitis (EAE), an MS model, confirmed elevated *Irg1* levels in myeloid, CD4, and B cells in the EAE group raising the concern if *Irg*1 is detrimental or protective. *Irg1* knockout (KO) mice exhibited severe EAE disease, increased mononuclear cell infiltration, and increased levels of triple-positive CD4+ T cells expressing IL17a, GM-CSF, and IFNγ. A lack of *Irg1* in macrophages elevates Class II expression, promoting the polarization of myelin-primed CD4+ T cells into pathogenic Th17 cells via the NLRP3/IL-1β axis. Adoptive transfer in Rag-1 KO and single-cell RNA sequencing highlighted the crucial role of *Irg1* in shaping pathogenic Th17 cells. Moreover, bone marrow chimeras revealed that immune cells lacking *Irg1* maintained pathogenic and inflammatory phenotypes, suggesting its protective role in autoimmune diseases, including MS.

**Significance:** Immunoresponsive gene 1 (*Irg*1) was identified as a significantly elevated gene under inflammatory conditions through in vitro and in vivo models. Using global knockout mice, we identified *Irg*1 as a protective endogenous gene that negatively regulates pathogenic Th17 cells. Single-cell RNA sequencing of infiltrating cells during EAE revealed that *Irg*1 knockout enhanced the expression of pathogenic Th signatures in CD4+ T cells, indicating a robust proinflammatory environment. *Irg*1 negatively regulates IL-1β in macrophages, which is essential for the differentiation of pTh17 CD4+ T cells, potentially clarifying the exacerbation of EAE in knockout animals. Our study identified *Irg*1 as a negative regulator of both innate and adaptive immune responses in a CNS autoimmunity model.

## INTRODUCTION

Multiple sclerosis (MS) is an autoimmune disorder of the central nervous system that mostly affects young individuals. It is influenced primarily by autoreactive CD4+ T cells, which are responsible for the demyelination process. An imbalance in the delicate regulation of innate and adaptive immune responses occurs in the context of MS, leading to changes in the presentation of self-antigens to CD4+ T cells by antigen-presenting cells (APCs) (1, 2). In the initial phase of the disease, both dendritic cells and macrophages function as local antigen presenters for CD4+ T cells, facilitating the development of T cells toward Th1/Th17 states. These Th cells are characterized by the production of cytokines such as IFNγ, IL17a, and GMCSF. These effector T helper cells enter the central nervous system (CNS) and are subsequently stimulated by local antigen-presenting cells (APCs), such as microglia, resulting in demyelination (3, 4).

The production of few immuno-metabolites is strictly controlled and occurs exclusively under inflammatory conditions. One such metabolite, itaconate, an offshoot metabolite of the TCA cycle, is recognized for its antibacterial properties and is closely associated with bacterial infections such as *Mycobacterium tuberculosis*(5). Immune-responsive gene 1 (*Irg1*) is a mitochondrial enzyme that produces itaconate at high concentrations (milli-molar) during inflammatory conditions, primarily in myeloid lineage cells(6). Itaconate is recognized for its antibacterial activities and ability to suppress the immune system, suggesting that the regulation of immune cell activity is a significant feature of itaconate (7–9).

Itaconate has been found to decrease the oxygen consumption rate, and additionally, it inhibits the activity of succinate dehydrogenase (SDH), leading to increased levels of succinate in activated macrophages(10, 11). Furthermore, studies have demonstrated that *Irg1* acts as an intermediary between the innate and adaptive immune responses(12). Its absence exacerbates proinflammatory responses, tissue damage and the recruitment of inflammatory cells (8). Itaconate has been shown to provide protection against autoimmune disease by regulating neuroinflammation(13) and restoring the balance of T cells through metabolic and epigenetic reprogramming in experimental autoimmune encephalomyelitis (EAE)(14). Nevertheless, the impact of *Irg1* in regulating CNS autoimmunity has not been investigated. The current work aimed to examine the impact of *Irg1* on the progression of EAE and assess its potential in controlling T-cell differentiation by using *Irg1* global knockout (KO) mice. Our observations revealed that the loss of *Irg1* led to significant alterations in the natural defense system of the body, resulting in increased severity of EAE. We observed aberrant expression of autoreactive triple-positive CD4+ T cells expressing IL17, GMCSF and IFNγ in the central nervous system, suggesting that *Irg1* expression is a crucial regulator of CD4+ T-cell function. Furthermore, macrophages lacking *Irg1* presented an increased ability to present antigens and induced the polarization of Th17 cells through the NLRP3/IL-1β pathway. This study revealed *Irg1* as a specific target for therapeutic intervention in the treatment of MS.

## RESULTS

### Induction of itaconate and *Irg1* in immune cells and the spinal cord during EAE

Previously, we reported altered metabolic signatures in the blood of MS patients and chronic and relapsing-remitting mouse models of EAE via untargeted global metabolic profiling(15–19), documenting blood-based alterations in metabolism during MS or EAE disease (20). To examine whether metabolic genes are perturbed in brain glial cells under inflammatory conditions, we generated rat brain primary mixed glial cells, treated them with LPS/IFNγ (0.5 µg/ml and 20 ng/ml) for 8 hours and processed them for RNA-seq. Interestingly, our results revealed that *Irg1* was among the top ten genes with the greatest fold change (padj-6.30E-09, logFC-9.9) (**Fig. 1A**). We validated the levels of *Irg1* at various time points and found that LI induced the expression of *Irg*1 at 4 h, which peaked at 8 h and was reduced at 16 h of treatment (**Fig. 1B**). To replicate the *in vitro* finding of increased *Irg1* levels under inflammatory conditions, we examined Irg1 expression in the spinal cord of EAE. We observed that *Irg1* expression at the protein level was significantly greater in the spinal cord of EAE mice than in that of CFA control mice, which was reflected by immunohistochemistry (IHC) of the spinal cord sections (**Fig. 1C-D**). Since *Irg1* has been reported to be increased in immune cells during infection or inflammation, we examined Irg1 expression in various cell types, including CD11b, monocyte, CD4 and B cells, during EAE. The expression of *Irg1* in these isolated immune cells was greater in the EAE group than in the CFA control group, which was further validated by quantitative PCR (**Fig. 1E-F**). Interestingly, we also observed significant upregulation of *Irg1* at both the mRNA and protein levels in CD4+ T cells from EAE mice compared with those from control mice, which was previously reported to involve *Irg-1* expression in myeloid cells(12, 21). These sets of experiments suggest that itaconate levels are increased as a result of increased *Irg1* expression in the CNS during EAE, raising the question of whether elevated Irg1 levels are protective or pathogenic.

**Figure 1:**
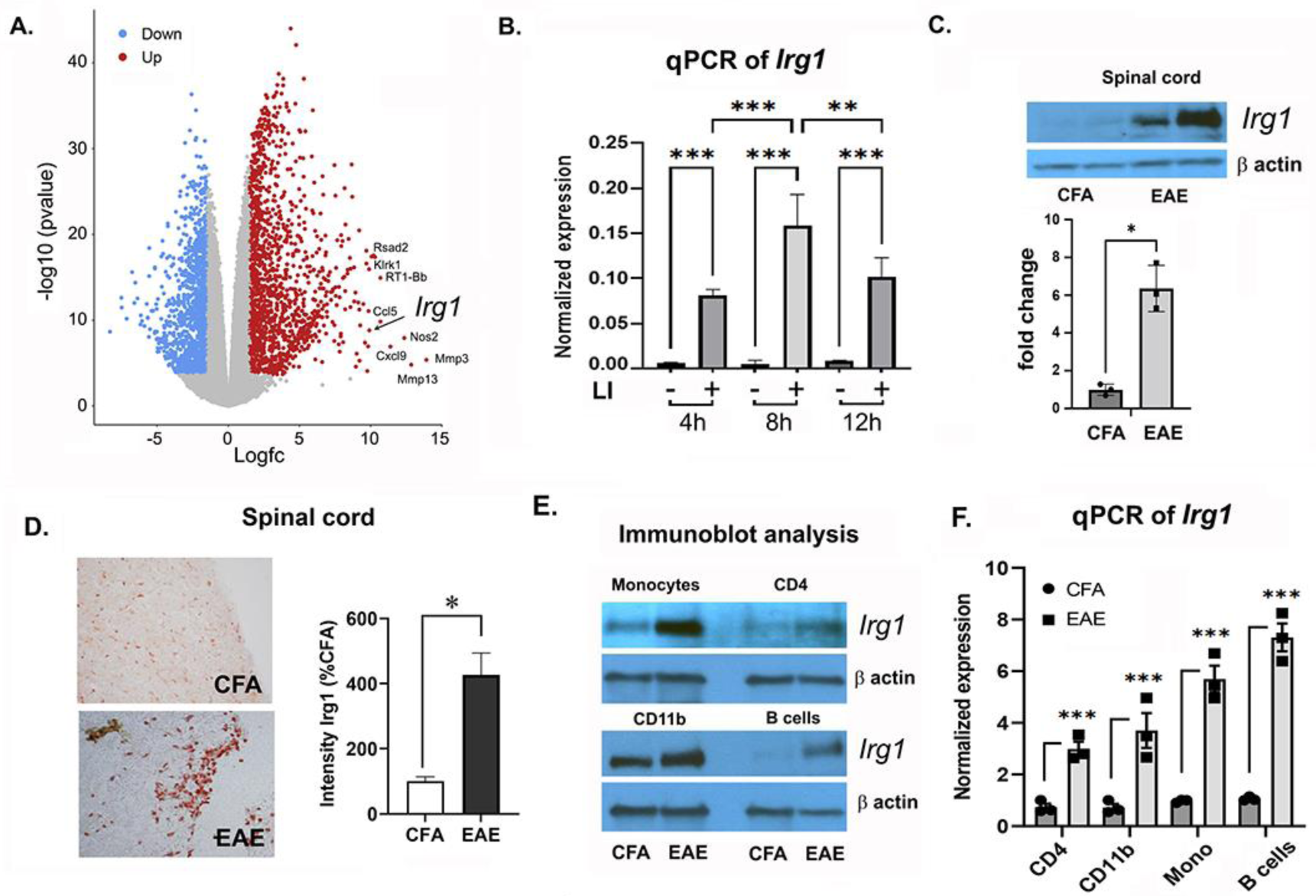
*Irg1* expression is induced in the spinal cords of EAE mice. **A.** Volcano plot showing the RNA-seq results showing significantly upregulated (red dots) and downregulated genes (blue dots) after 8 h of LI treatment in comparison with those of untreated rat brain mixed glia, with the top 10 dysregulated genes labeled in black (Padj <0.0005 & logFC >1.5) (N=4). **B.** *Irg1* expression was examined in rat brain mixed glia treated with or without LI for 4 hrs, 8 hrs and 16 hrs (n=3). (**\*\*\*** p >10^−6^). **C.** qPCR and immunoblot data revealed the upregulation of *Irg1* (normalized to β-actin) in the spinal cord of EAE mice. Data are representative of three independent experiments. **D.** Representative images of the spinal cord (CFA vs EAE) stained with *Irg1*. Bar graph representing the quantification of *Irg1* expression intensity as a percentage of CFA from a total of three independent experiments. **E.** Quantitative analysis of *Irg1* expression in monocytes, CD4+ cells, CD11b^+^ macrophages and B cells, presented as the relative protein expression normalized to that of the housekeeping control β-actin and expressed as the fold change compared with that of CFA. **F.** The results from the qPCR analysis of *Irg1* expression in monocytes, CD4+ cells, CD11b^+^ macrophages and B cells from CFA and EAE mice normalized to that of β-actin, and the data are representative of three independent experiments. The data presented in the bar graph are the means ± SDs. *p < 0.05, **p < 0.01, and ***p < 0.001 compared with the LI or EAE group using Student’s t test or one-way ANOVA.

### Loss of *Irg1* led to severe EAE and the activation of immune cells

To evaluate the significance of increased *Irg1* expression in the spinal cord in EAE, we induced EAE in global *Irg1* knockout (KO) mice via MOG35-55 (200 µg/ml) as described previously. We observed a significant difference in the clinical score and cumulative score between *Irg1*-KO mice and Wt mice (**Fig. 2A-B**). Upon examination of the antigen-specific response, spleen/lymph node *Irg1* KO cells produced significantly increased levels of IFNγ, IL17a and GM-CSF without affecting IL4 upon MOG35-55 stimulation (**Fig 2C**). Furthermore, histopathology analysis revealed greater inflammatory cell infiltration and demyelination in the spinal cord of *Irg1*-KO mice than in the spinal cord of Wt mice, as revealed by H&E and LFB staining, respectively (**Fig. 2D**), which correlates with the severity of the clinical score in *Irg1*-KO mice. The infiltration of mononuclear cells, including myeloid cells and T cells that influence the CNS environment, is detrimental to oligodendrocytes and neurons, leading to demyelination in MS and EAE (22). Similar to the H&E staining data, we also observed a significant increase in infiltrating mononuclear cells (CD45^hi^ populations) in the CNS tissue of *Irg1-*KO mice with EAE compared with that in Wt EAE mice (**Fig. 2E**). Immune profiling followed by tSNE projection of the CNS infiltrates at the peak of the disease revealed a greater number of CD4+ T cells in *Irg1*-KO mice than in WT mice. Hence, we quantified the absolute number of infiltrating CD4+ T cells (CD45^+^CD3^+^CD4^+^) and found that *Irg1*-KO mice with EAE presented ∼3-fold more CD4+ T cells than did WT mice (**Fig 2G**). CNS infiltrates in *Irg1* KO mice presented higher levels of Th1- and Th17-expressing double-positive CD4+ T cells (expressing IL17 and GM-CSF) and triple-positive CD4+ T cells expressing IL17, GM-CSF and IFNγ than WT infiltrating cells did (**Fig 2I**). Since myeloid cells are important for disease induction and disease severity due to their role in differentiating antigen-specific Th cells in the peripheral lymphatic organ and within the CNS (2), we profiled myeloid cells in Wt and KO CNS tissues. Myeloid cell profiling via tSNE projection of CNS infiltrates revealed increased infiltration of CD11b+ myeloid cells, monocytes/macrophages and dendritic cells (DCs) (**Fig 2F**, **2J**). Similar observations were made in the spleen, as significant increases in two different macrophage populations, CD11b^low^F4/80^+^ (red pulp macrophages)(23) and CD11b^med^F4/80^+^ (migrated macrophages), were observed (**Supplementary Fig. 1-2**).

**Figure 2:**
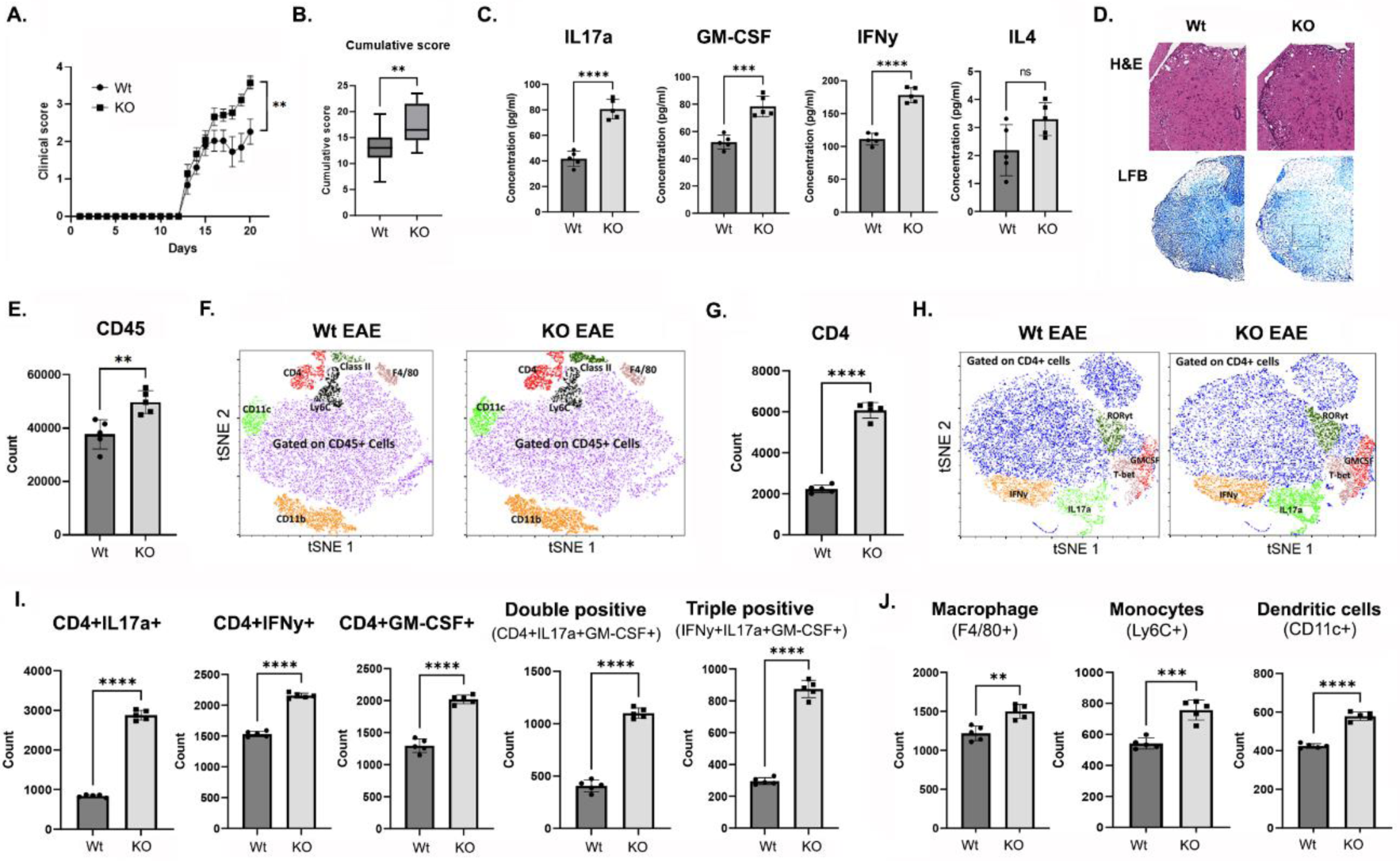
Irg1-KO mice exhibited more severe disease. **A.** Wt and *Irg1*-KO female mice were immunized with MOG35-55 (200 µg/mouse) on day 0 in CFA, and PT was given on days 0 and 2 of immunization. Clinical scores were recorded until 22 d (n = 22). **B.** Bar graph representing the cumulative score of disease severity in Wt and *Irg1*-KO mice (n=22). **C.** Recall response was assessed via ELISA of spleen and LN samples isolated from Wt & *Irg1-*KO mice (n=5) subjected to MOG (20 µg/ml) treatment. Samples were collected at 72 hr, and the levels of the cytokines IL17a, GM-CSF, IFNγ and IL4 were measured, with error bars representing the S.D. (n=5). **D.** Representative images showing histopathological changes in the spinal cord tissues of EAE mice in the Wt and *Irg1*-KO groups at 22 days postimmunization. Sections were stained with hematoxylin and eosin (H&E) and Luxol fast blue (LFB) to visualize cell infiltration and changes in myelin content. **E.** Flow cytometry analysis revealed the total number of CD45+ cells in the CNSs of Wt and *Irg1*-KO mice (n=5). **F.** Flow cytometry t-SNE plot gated on CD45^+^ cells in BILs from Wt and *Irg1-*KO mice demonstrating myeloid cell populations; CD11b^+^, F4/80^+^, Ly6C^+^, and CD11c^+^ populations. **G.** Flow cytometry analysis showing CD4+ populations in the CNSs of Wt and *Irg1*-KO mice (n=5). **H.** Representative flow cytometry t-SNE plot of BILs gated on CD4^+^ cells demonstrating the IL17a, IFNg, T-bet, RORγt and GM-CSF populations. **I.** Bar graph showing CD4+ cells expressing the total positive population of IL17a, IFNg, and GMCSF, as well as double-positive (IL17-A+IFNγ) and triple-positive (IL17-A+IFNγ+GMCSF) cells in Wt (n=5) and *Irg1* KO (n=5) strains. **J.** Bar graph showing the total positive population of infiltrating macrophages, monocytes and dendritic cells. The data presented in the bar graph are the means ± SDs. *p < 0.05, **p < 0.01, and ***p < 0.001 compared with the *Irg*1 KO group using Student’s t test or one-way ANOVA.

### Single-cell RNA sequencing of *Irg*1 KO mice with EAE revealed hyper infiltration of CD4+ T cells

To determine the effect of the loss of *Irg*1 on CNS inflammation, we analyzed brain and spinal cord cellular heterogeneity and the transcriptome at single-cell resolution by performing PIP-seq in *Irg1*-KO and WT mice with EAE on day 8 postimmunization (**Fig. 3A**). At a resolution of 0.25, we identified 11 transcriptionally distinct cell clusters as visualized by Uniform Manifold Approximation and Projection (UMAP) (**Fig. 3B**). On the basis of the expression of known marker genes, distinct cell clusters, including CD4+ and CD8+ T cells, macrophages, microglia, and neutrophils, as well as oligodendrocytes, were identified (**Fig 3C-D**). Compared with WT *Irg1* KO mice, *Irg*1 KO mice presented drastic differences in cell numbers (**Fig. 3E**) and gene expression (**Fig. 3F-H**). In particular, compared with WT mice with EAE, *Irg1* KO mice presented significant increases in CD4+ T cells, CD8+ T cells, and Ly6C+ macrophages and substantial reductions in oligodendrocyte populations. Moreover, *Irg1*-KO mice presented increased total infiltration of immune cells into the spinal cord, as evidenced by the increased number of CD45+ (a pan leukocyte marker) cells (**Fig. 3F**). Furthermore, *Irg1* KO upregulated the expression of pathogenic Th signatures such as IFNγ, IL-17a and GM-CSF in CD4+ T cells (**Fig. 3G**), suggesting a strong proinflammatory milieu, which might explain the exacerbation of EAE in the KO mice. The *Irg1*-KO mice presented significantly increased antigen presentation and inflammatory signaling, as demonstrated by elevated numbers of infiltrating macrophages expressing MHC class II and IL-1β (**Fig. 3H**).

**Figure 3:**
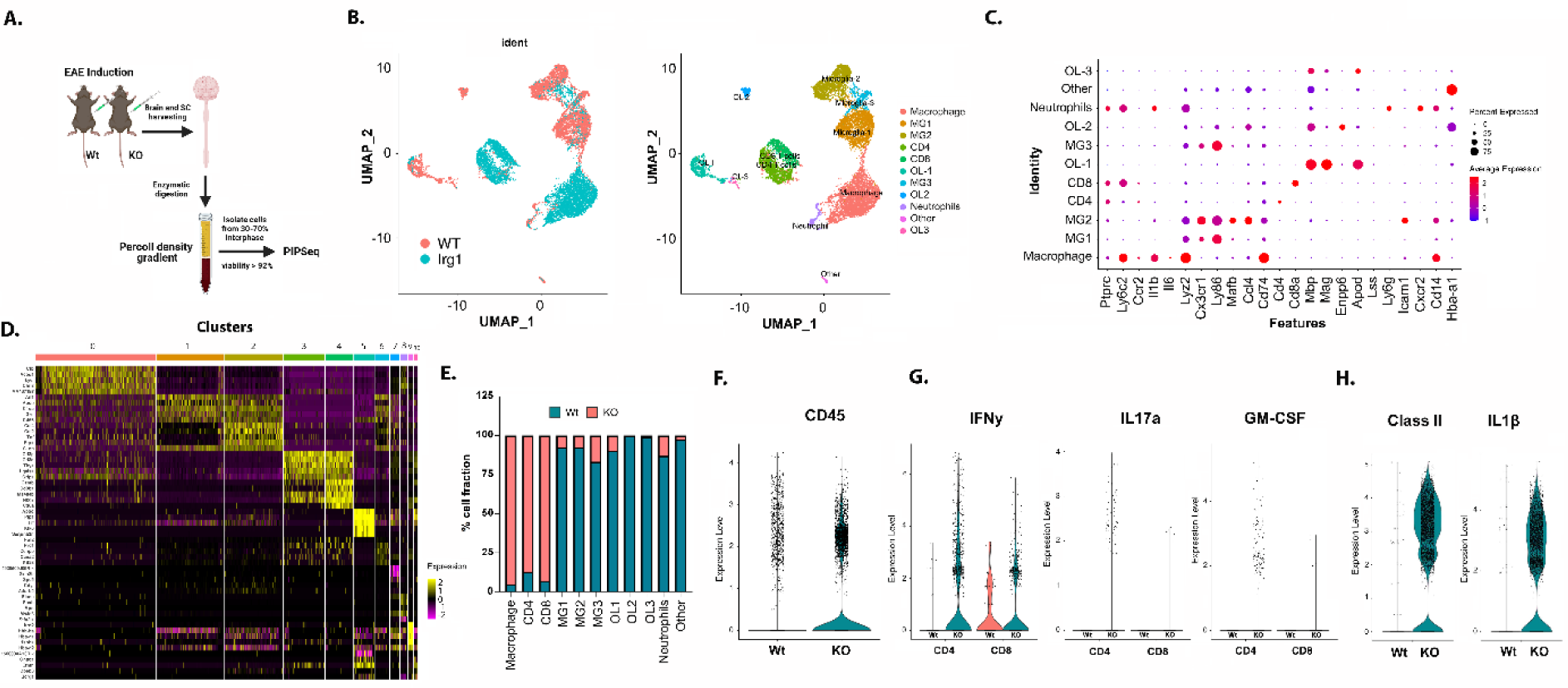
scRNA-seq of CNS tissues from *Irg*1 KO mice revealed increased infiltration of immune cells during EAE. **A.** A schematic representation of scRNA sequencing conducted on the spinal cord from EAE-immunized wild-type and Irg1Ko mice via PIP-Seq. A single-cell suspension of the spinal cord was prepared via Percoll density gradient, and total RNA was isolated and subjected to high-throughput sequencing via PIP-Seq. **B.** Sc-RNA-seq clusters represented by UMAP, with the first plot showing differences in the cell clusters between the two groups, WT (blue) and Irg1 KO (salmon), and the second plot showing the cell clusters, where each color represents a different cell type with the color key on the right. **C**. Dot plots showing signature gene expression across the 11 cell clusters. The size of the dots represents the proportion of cells expressing a particular marker, and the color spectrum indicates the mean expression levels of the markers. **D.** Heatmap showing the expression of the top 5 marker genes across the 11 cell clusters. **E.** Bar graph showing the percentages of all 11 cell types. **F.** Violin plots of normalized expression levels of the CD45 gene in WT and Irg1 ko plants grouped by conditions. **G.** Violin plots of normalized expression levels of the IFNγ, Il17a, and GMCSF genes in Wt and Irg1-KO CD4 and CD8 T cells. **H.** Violin plots of the normalized expression levels of the MHC-II and IL-1β genes in Wt and Irg1-KO macrophages.

### CD4 cells lacking *Irg1* are highly pathogenic in nature

As mentioned above, we observed that CD4+ *Irg1* exacerbated the expression of inflammatory cytokines (IL17a, GM-CSF and IFNγ); therefore, we next aimed to determine whether antigen-specific CD4+ cells lacking *Irg1* are encephalitogenic in nature. We performed adoptive transfer (Adt) of MOG35-55-primed CD4 cells lacking *Irg1* and Wt into Wt B6 mice. For this purpose, we immunized Wt and *Irg1*-KO mice with MOG35-55, and after 10 days, spleen/LN cells were stimulated with antigen in the presence of anti-IFNγ (10 µg/ml) and IL12p70 (20 ng/ml). After 72 h, we found that CD4 T cells lacking *Irg1* produced higher levels of IL17a and GM-CSF than did Wt CD4 T cells (**Fig. 4A**). Upon adoptive transfer of these cells into Wt mice, we found that compared with Wt mice receiving MOG-primed Wt CD4 cells, Wt mice receiving MOG-primed CD4 cells lacking *Irg*1 presented severe EAE pathogenicity (**Fig. 4B**). Furthermore, Rag1 mice, which are deficient in resident T and B cells, also showed severe EAE pathogenicity in response to Adt of MOG35-55-primed CD4 T cells lacking *Irg1* (KO->Rag1) compared with Wt CD4 cells (Wt->Rag1) (**Fig. 4C**). The severity of clinical symptoms in the KO->Rag1 group was reflected by significantly greater infiltration of CD4+ T cells in the CNS with the Th17 phenotype. However, we did not observe a significant but trending increase in the population of CD4-expressing IFN-γ^+^ and GMCSF^+^ populations, with no change in Th2 cells in the CNS in the KO->Rag1 group compared with the Wt->Rag1 group (**Fig. 4D-H**). More importantly, there was an exacerbation of double (CD4^+^IL17^+^GMCSF^+^ and CD4^+^IL17^+^IFN-γ^+^)- and triple (CD4^+^IL17^+^GMCSF^+^IFNγ^+^)^-^ positive cells in the CNSs of the KO->Rag1 to Wt->Rag1 groups with EAE (**Fig. 4I-K**). These sets of data clearly suggest that CD4 cells lacking *Irg1* are highly pathogenic in nature and that the expression of *Irg1* may be one of the crucial regulators of CD4-mediated pathogenicity in autoimmune diseases such as EAE.

**Figure 4:**
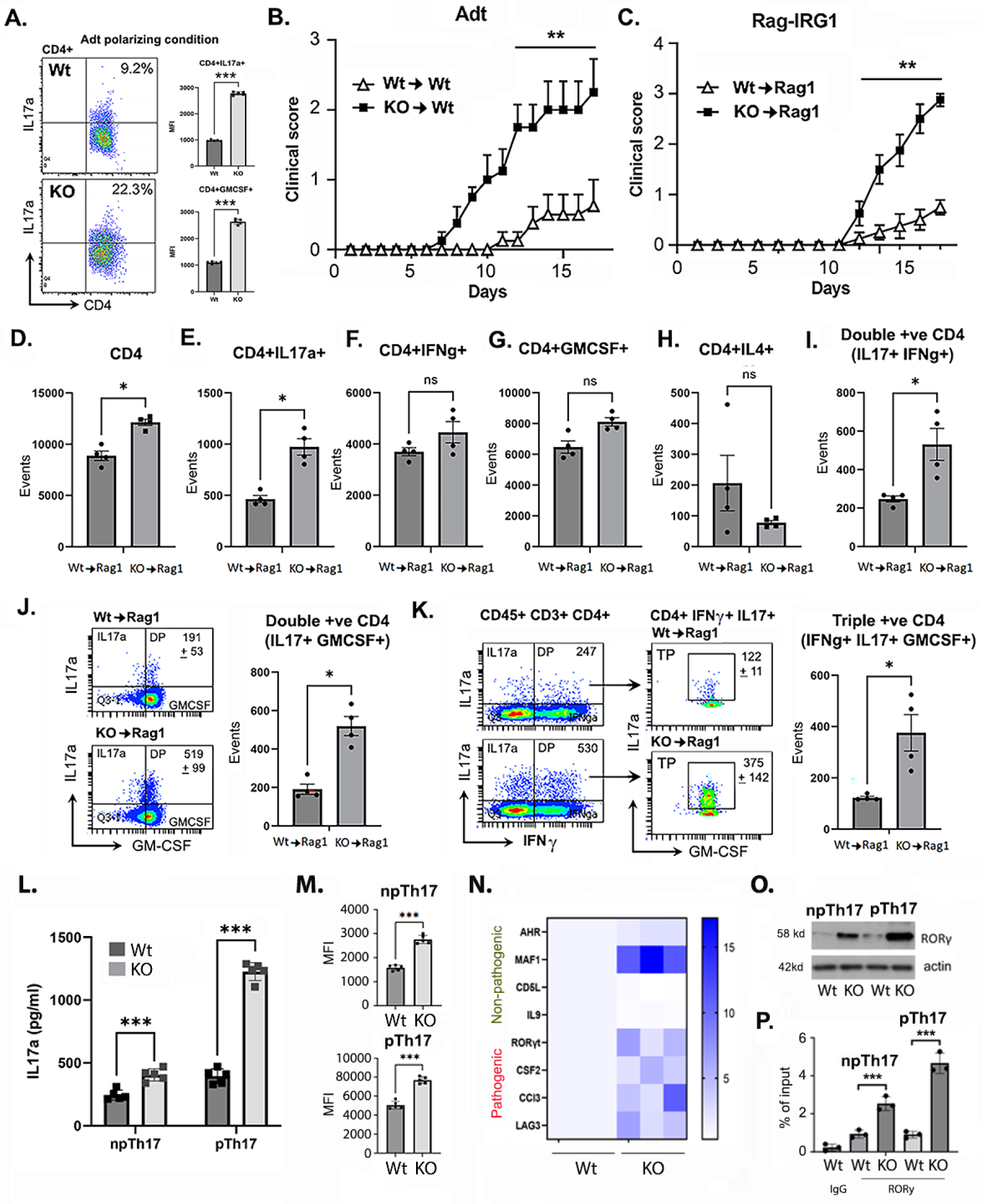
*Irg1* KO CD4 cells are highly pathogenic in nature. **A.** Quantification of CD4^+^IL17a^+^ cells after CD4^+^ cells from Wt and *Irg1*-KO mice were adoptively transferred into B6 mice (n=5). **B.** Clinical scores were obtained until 17 d after B6 mice were immunized with MOG35-55 (200 µg/mouse) (n=10). **C.** Clinical scores were obtained for Rag1 mice after adoptive transfer of CD4^+^ cells from Wt and *Irg1*-KO mice, followed by immunization with MOG35-55 (200 µg/mouse) and PT (n=10). The bar graph represents the positive populations of (D) total CD4^+^ cells (**E**), CD4^+^IL17a^+^ cells (**F**), CD4^+^ IFNγ^+^ cells (**G**), CD4^+^ GMCSF^+^ cells (**H**), CD4^+^ IL4^+^ cells and (**I**) double-positive (IL17a^+^IFNγ^+^) in Wt and *Irg1* KO CD4^+^T cells received Rag1 mice respectively (N=4). **J**. Flow cytometry plot (left) gated on CD4+ cells showing double-positive (IL17a^+^GMCSF^+^) populations in Rag1 mice, which are presented as a bar graph (right) (N=4). **K.** Flow cytometry plot (left) gated on CD4+ cells showing triple-positive (IL17a^+^ IFNγ^+^ GMCSF^+^) populations in Rag1 mice, which are presented as a bar graph (right) (N=4). **L.** ELISA analysis of the levels of the cytokine IL17a in pathogenic and nonpathogenic Th17 cells isolated from Wt and *Irg1*KO mice (N=5). **M.** Representative flow cytometry plots (left) of IL17a^+^ populations in npTh17 and pTh17 cells and data are presented as a bar graph (right) showing the number of IL17a^+^ cells (MFI) in npTh17 and pTh17 cells (N=5). **N.** Heatmap showing the mRNA expression of Th17 pathogenic and nonpathogenic signature genes in Wt and KO mice (N=3). **O.** Immunoblot data of RORγt (normalized to β-actin) in npTh17 and pTh17 cells from Wt and *Irg*1-KO mice. The data are representative of three independent experiments. **P.** A ChIP assay was performed to determine the binding of RORγt to the IL17a promoter. The data are presented as the percentage of RORγt binding to the IL17 promoter with respect to the total input. NS not significant, *p < 0.05, **p < 0.01, and ***p < 0.001 compared with the Irg1-KO group using Student’s t test or one-way ANOVA.

### *Irg1* KO potentiates pathogenic Th17 subsets upon EAE induction

The Th17 subset is critical to EAE progression and is known to include two distinct subsets depending on their pathogenic inflammatory profile (24). One subset is known as the pathogenic Th17 (pTh17) subset, which produces high levels of inflammatory mediators, such as IL17a and GMCSF, and induces EAE disease (25–27), whereas the second subset is the nonpathogenic Th17 (npTh17) subset, which has a very mild inflammatory profile (28, 29). Since we observed that CD4 T cells lacking *Irg1* presented increased Th17 numbers with encephalitogenic properties, we examined whether CD4 T cells lacking *Irg1* expression were predisposed to the pathogenic Th17 phenotype. For this purpose, we cultured naive CD4 T cells from Wt and *Irg1* KO mice and cultured them under npTh17 (TGFβ and IL6) and pTh17 (TGFβ, IL6, IL23 and IL1β) conditions as described previously (24). We observed that CD4 cells lacking *Irg1* produced significantly higher levels of IL17a under both pathogenic and nonpathogenic conditions (**Fig. 4L**). This observation was further supported by flow cytometry intracellular staining, which revealed a greater number of IL17-producing Th17 *Irg1*-KO CD4+ T cells under both conditions (**Fig. 4M**). A set of differentially expressed genes specific to the pTh17 and npTh17 phenotypes (24) were examined, and we observed significantly greater upregulation of genes associated with pathogenic Th17 cells, including RORγt, CSF2, CCl3 and LAG3, in CD4 cells lacking *Irg1* than in Wt CD4 cells, with downregulation of genes associated with nonpathogenic Th17 cells, such as CD5L and IL-9 (**Fig. 4N**). Although the upregulation of some genes associated with nonpathogenic Th17 cells, such as AHR and MAF-1, in CD4 cells lacking *Irg1*, the AHR and MAF-1 genes are reportedly associated with the proliferation of Th17 cells. These results suggest that *Irg1* negatively regulates both the pathogenicity and proliferation of Th17 subsets. Th17 differentiation is programmed by the nuclear factor retinoid-related orphan receptor-gamma (RORγt)(30, 31). As depicted in **Figure 4N & O**, CD4 T cells lacking *Irg1* expression expressed a greater level of RORγt than Wt CD4 T cells under npTh17 and pTh17 conditions. To clarify the detailed mechanism, we used a ChIP assay to establish whether increased levels of RORγt in CD4+ T cells lacking *Irg1* are responsible for the pathogenic Th17 phenotype. We found that the recruitment of RORγt to the promoter of IL17a in Irg1-deficient CD4+ T cells was significantly greater than that in WT CD4+ T cells **(Fig. 4O-P)**. Rabbit IgG was used as a negative control, and no PCR product could be detected, suggesting the specificity of the CHIP assay. This set of data clearly suggests that *Irg1* negatively regulates RORγt expression and its recruitment to influence Th17 cell differentiation and pathogenicity.

### *Irg1*-deficient macrophages are proinflammatory in nature

Myeloid cells, mainly monocytes/macrophages, are the major effectors of demyelination in both MS and EAE(32–34). A study reported that macrophages lacking *Irg1* presented a hyperinflammatory response after Mycobacterium infection(8). Here, we examined the status of proinflammatory cytokine expression in Wt and *Irg1*-KO macrophages. We found that compared with Wt macrophages, macrophages lacking *Irg1* showed increment in the level of TNFα, IL6, IL1β and MCP1 upon LPS stimulation.) (**Supplementary Fig. 3A**). Similar results were observed when splenic macrophages were isolated from the spleens (anti-F4/80 microbeads, Miltenyi Biotec) of Wt and *Irg1*-KO EAE mice, and their expression was examined via quantitative PCR (**Supplementary Fig. 3B**). Furthermore, qPCR revealed that *Irg1* KO BMDMs showed significant upregulation of iNOS expression, whereas a differential effect was observed for anti-inflammatory-associated genes, such as an increase in EGR2 and a decrease in Arg1, compared with those in Wt BMDMs, with no change in CD206 (**Supplementary Fig. 3C**). Macrophages are known for their phenotypic transitions; therefore, to elucidate the impact of *Irg1* loss on their phenotypic transitions, BMDMs were stimulated with IFNγ (50 ng/ml), and phenotypic characterization was performed via flow cytometry. Compared with Wt BMDMs, *Irg1* KO BMDMs were significantly pushed toward a proinflammatory state as characterized by F4/80^+^CD86^+^ and F4/80+Class II+ phenotypes (**Supplementary Fig. 4A**). The observed increase in proinflammatory cytokines (IL6 and IL1β) is well known to potentiate and maintain the Th17 phenotype(35), suggesting that the increased expression of IL17 in *Irg1*-KO EAE mice could be due to increased levels of these cytokines.

### Loss of *Irg1* resulted in increased antigen presentation by macrophages

To further confirm that infiltrated *Irg1* KO macrophages are proinflammatory in the CNS, we profiled infiltrated macrophages expressing Class II in *Irg1* KO and WT mice with EAE. We found that the expression of Class II was significantly greater in infiltrated macrophages (CD11b^+^F4/80^+^ cells) in Irg1 KO than Wt macrophages (**Fig. 5A & Supplementary Fig 4B**). We also examined class II expressions on other infiltrating myeloid cells in Wt and *Irg1*-KO mice with EAE. The tSNE projection of Class II+CD45-infiltrated myeloid cells revealed that all myeloid cells, including CD11b+ cells, Ly6C+ monocytes, CD11c+ cells and macrophages (F4/80), presented higher Class II expression in *Irg1* KO mice than in Wt mice (**Fig. 5B-C**). We further characterized the immune profile-specific cell population and revealed that infiltrating inflammatory dendritic cells (DCs; CD45+CD11b-CD11c+Class II+), monocytic DCs (CD45+CD11b+CD11c+), monocytes (CD45+CD11b+Ly6G^-^Ly6C+) and brain-resident microglia (CD45^lo^CD11b+) were more highly expressed in Class II *Irg1*-KO mice than in Wt EAE mice (**Fig. 5D-G**), suggesting their potential ability to present antigens to myelin-specific CD4+ T cells. Additionally, a similar trend was found in splenic macrophages (**Supplementary Fig. 1-2**).

**Figure 5:**
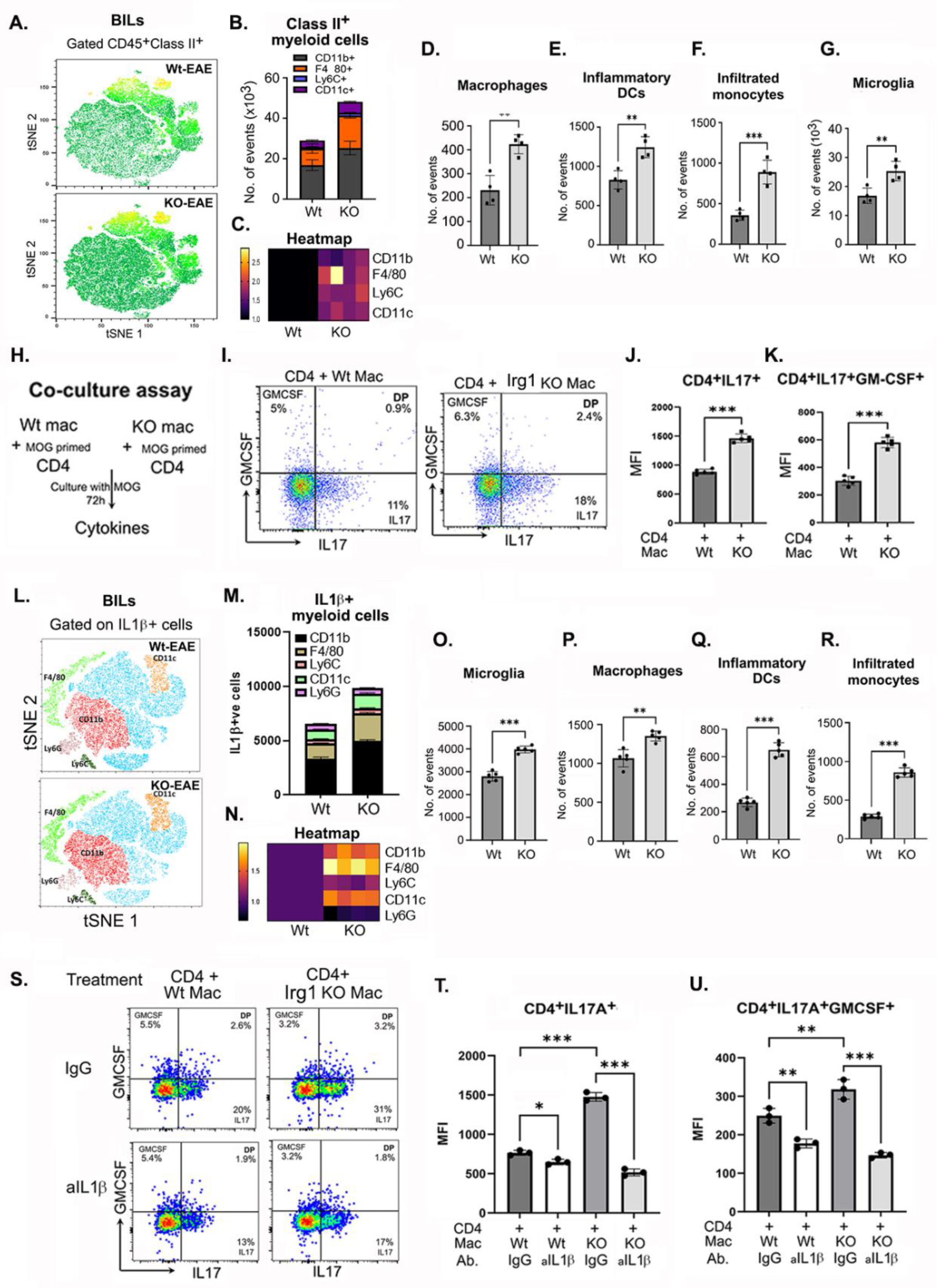
IL-1β plays a key role in the antigen presentation of *Irg1* KO macrophages and is more potent in polarizing MOG-primed CD4 T cells toward pTh17 cells. **A.** Representative flow cytometry t-SNE plot of BILs gated on CD45^+^ cells demonstrating Class II^+^ populations in Wt and *Irg1*-KO mice and (**B-C.**) The bar graph shows populations of Class II-expressing myeloid cells, namely, CD11b^+^, F4/80^+^, CD11c^+^ and Ly6C^+^ cells (N=5). **D-G.** The bar graph represents populations of Class II infiltrating macrophages (CD45^hi+^CD11b^+^F4/80^+^), inflammatory DCs (CD45^+^CD11c^+^Class II^+^), infiltrated monocytes (CD45^+^CD11b^+^Ly6C^+^Class II^+^) and microglia (CD45^low^CD11b^+^Class II^+^) (N=5). **H.** Schematic overview of the coculture study using Wt CD4 cells with Wt and *Irg1*-KO macrophages. **I-K.** Flow cytometry data gated on CD4^+^ cells demonstrating the populations of double-positive (IL17a^+^GM-CSF^+^) upon coculture of WT CD4 T cells with WT and *Irg1-*KO macrophages and (J-K) MFI values of positive populations of IL17a^+^, GMCSF^+^ and double-positive (IL17a^+^GM-CSF^+^) T cells are presented as a bar graph. (N=5). **L.** Representative flow cytometry t-SNE plot gated on ^IL-1β+^ cells demonstrating populations of IL-1β-producing various myeloid cells; CD11b^+^, F4/80^+^, CD11c^+^, Ly6C^+^ and Ly6G^+^ cells. **M.** Bar graphs represent the positive population of myeloid cells producing IL-1β in WT and *Irg1-*KO mice. **N.** Heatmap showing the positive population of myeloid cells producing IL-1β in WT and *Irg1-*KO mice (N=4). **O-R.** The bar graph represents the total number of IL-1β+ve-expressing microglia (CD45^low^CD11b^+^), infiltrated macrophages (CD45^hi+^CD11b^+^F4/80^+^), inflammatory DCs (CD45^+^CD11c^+^Class II^+^) and monocytes (CD45^+^Ly6C^+^Ly6G^-^). **I.** Flow cytometry plot demonstrating populations of IL17a^+^ and IL17a^+^GMCSF^+^ CD4^+^ cells upon coculture with Wt and *Irg1*KO macrophages in the presence or absence of αIL-1β (10 ng/ml). **T-U.** A population of IL17a^+^ and double-positive (IL17a^+^GMCSF^+^) CD4+ cells is presented as a bar graph after myelin-specific CD4+ T cells were cocultured with Wt and *Irg1* KO macrophages in the presence or absence of αIL1β (n=3). The data presented here are the means ± SDs. NS, not significant; *p<0.05, **p<0.001, ***p<0.0001 compared with *Irg1* KO macrophages cocultured with CD4 cells; Student’s t test, one-way ANOVA.

To test this hypothesis, we generated macrophages from the bone marrow of both Wt and *Irg1*-KO mice, cultured them with 2D2 CD4+ T cells in the presence of MOG35-55 and examined the production of proinflammatory cytokines via ELISA and phenotypic characterization via flow cytometry (**Fig. 5H**). We observed that 2D2 CD4 T cells cocultured with *Irg1*-KO macrophages presented significantly higher levels of IL17a, GM-CSF and IFNγ (**Supplementary Fig. 4C**) than Wt macrophages cocultured with 2D2 CD4 T cells. Similar observations were made when CD4+ cells were subjected to intracellular staining for IL17a, GM-CSF and double-positive (CD4+IL17a+GM-CSF+) (**Fig. 5I-K**). These sets of experiments clearly demonstrate that *the* loss of *Irg1* potentiates the ability of macrophages to efficiently present antigens to myelin-specific CD4+ T cells and polarizes them into Th17 cells expressing GM-CSF.

### Monocytes/macrophages are the major producers of IL-1β in *Irg1*-KO mice

Myeloid cells are the main source of IL-1β, which is a key player in polarizing pathogenic Th17 cells (36, 37). Since *Irg1*-deficient macrophages produce increased levels of IL-1β and other studies have reported similar outcomes, we postulated that IL-1β drives the pathogenic Th17 phenotype in *Irg1*-KO mice. First, we determined which myeloid population is the major contributor of IL-1β in the brain-filtrating cells (BILs) of Wt and *Irg1*-KO mice via t-distributed stochastic neighbor embedding (tSNE) plots and found that macrophages (F4/80+ population) and CD11c+ cells are major producers of IL-1β in the BILs of Wt and *Irg1*-KO mice (**Fig. 5L‒N**). Compared with Wt macrophages, *Irg1*-deficient macrophages produced significantly higher levels of IL-1β (**∼2-fold**), as reflected in the bar graph and heatmap (**Fig. 5M-N**). Since microglia are known as brain-resident macrophages and play a key role in neuroinflammation(38, 39), we also examined their contribution to the production of IL-1β in *Irg1*-KO mice with EAE. We observed that *Irg1*-deficient microglia produced significantly more IL-1β than did Wt microglia (**Fig. 5O**). Immunophenotyping revealed that other myeloid populations, including infiltrating inflammatory dendritic cells (DCs; CD45+CD11b-CD11c+Class II+), classical monocytes (CD11b^+^CD45^+^Ly6C^high^) and infiltrated macrophages (CD11b^+^F4/80^+^), were significantly more likely to produce IL-1β in the CNS of *Irg1* KO mice than in those of Wt mice with EAE (**Fig. 5P-R**). Additionally, both red pulp splenic macrophages and migrated macrophages significantly increased the production of IL-1β in *Irg1*-KO mice compared with Wt mice with EAE (**Supplementary Fig. 1,2 & 5**). On the basis of these observations, we concluded that a lack of *Irg1* led to the upregulation of macrophage/monocyte production of proinflammatory cytokines, mainly IL-1β, possibly pushing myelin-specific CD4 T cells toward pathogenic Th17 cells.

### *Irg1* negatively regulates the inflammasome in macrophages during EAE

IL-1β is an NLRP3 inflammasome-dependent cytokine, and NLRP3 has been reported to play a critical role in the development of EAE by regulating Th17 (40) and chemotactic immune cell migration (41). Since we detected increased levels of IL-1β in *Irg1* KO mice with EAE and *Irg1* negatively regulates NLRP3 activity(42, 43) in macrophages, we examined NLRP3 inflammasome components. For this purpose, Wt and *Irg1*-deficient macrophages were treated with LPS for 6 h, and the cell lysates were processed for the expression of IL-1β, NLRP3, ASC and GSDMD via immunoblot analysis. As depicted in **Supplementary Fig. 6A**, *Irg1*-deficient macrophages produced higher levels of pro-IL-1β and cleaved IL-1β than did WT macrophages under LPS stimulation. Like IL-1β-deficient macrophages, Irg1-deficient macrophages expressed higher levels of NLRP3 and GSDMD without any change in the level of ASC, suggesting that the level of the NLRP3 inflammasome is greater. Higher levels of IL-1β and NLRP3 were further validated via qPCR. As depicted in **Supplementary Fig. 6B**, stimulation of Wt macrophages with LPS induced the expression of IL-1β and NLRP3, which were significantly increased in *Irg1*-deficient macrophages under LPS stimulation, suggesting that *Irg1* negatively regulates NLRP3 and IL-1β in macrophages.

We previously demonstrated that *Irg1*-KO mice presented greater numbers of pTh17 CD4 cells than did WT mice with EAE, in which IL-1β expression plays an important role in pTh17 differentiation(36, 37); therefore, to further understand the importance of IL-1β in *Irg1*-mediated Th17 pathogenicity, we cocultured *Irg1*-KO macrophages with myelin-specific CD4 cells with either IgG or an αIL-1β neutralizing antibody (10 µg/ml) (**Fig. 5S**). We observed that, compared with IgG alone, neutralization of IL1β significantly diminished the population of IL17-A^+^, GMCSF^+^, and double-positive populations (IL17-A^+^GMCSF^+^) (**Fig. 5S-U**), whereas *Irg1*-KO macrophages were cocultured with antigen-specific CD4+ T cells. Our observations suggest that *Irg1* negatively regulates IL-1β in macrophages, which plays a critical role in differentiating pTh17 CD4+ T cells, resulting in severe EAE in *Irg1*-KO mice.

### Irg1-deficient immune cells maintain a proinflammatory nature in the CNS

Our data, both *in vivo* and *in vitro*, clearly demonstrated that *Irg1* loss significantly induced pathogenic Th17 cells and proinflammatory macrophages. Here, we aimed to investigate the impact of *Irg1* loss on immune cells in the same microenvironment. We generated chimeric mice with CD45.1 Wt and CD45.2 *Irg1* KO bone marrow via the busulfan method. After 6 weeks, when chimeras were established (**Supplementary Fig. 7**), EAE was induced via MOG35-55 as described above. At the peak of EAE disease, BILs were isolated from the brain and spinal cord via Percoll gradient, CD45.1 and CD45.2 cells were sorted via FACS, and RNA-seq was performed (**Fig. 6A‒ C**). A comparative analysis of the two groups revealed that 1,457 genes were significantly downregulated and that 1541 genes were significantly upregulated (**Fig. 6D**). We first confirmed the reduction in Irg1 expression in the CD45.2 group, along with its dependent genes, Nrf2 and Hmox1(44). We observed a significant reduction in Irg1, Nrf2 and Hmox1 expression in the CD45.2 group compared with the Cd45.1 group (**Fig. 6E**). We also observed significantly increased expression of CD45, suggesting increased infiltration of peripheral immune cells in the brains of *Irg1*-KO mice with EAE, which is supported by our flow cytometry data from BILs (**Fig. 2**, **3**).

**Figure 6:**
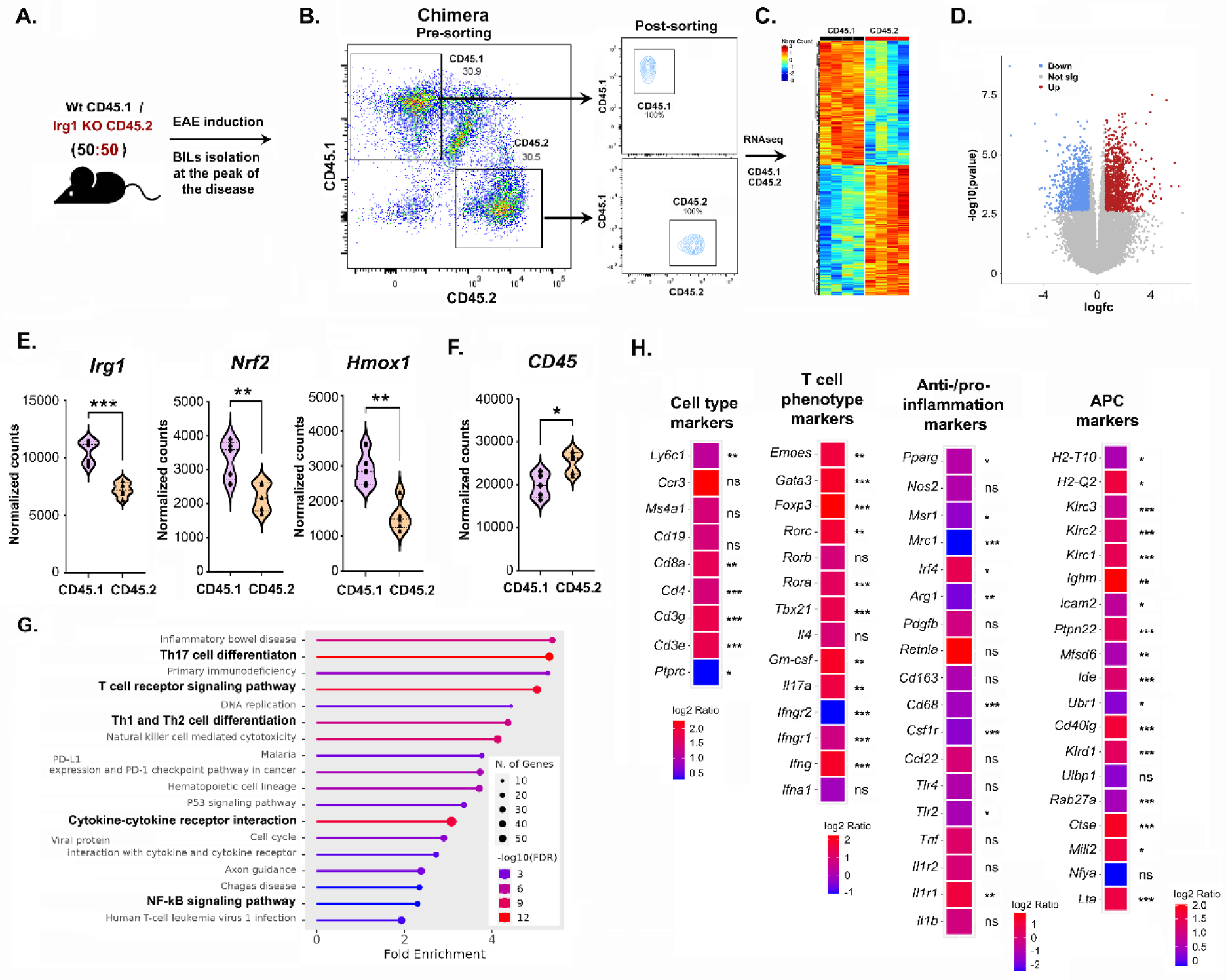
*Irg1* expression controls the proinflammatory and pathogenic signatures of infiltrating immune cells in the CNS. **A.** Chimera was generated via the use of 50:50 CD45.1 (Wt) and CD45.2 (*Irg1* KO) BMs utilizing Bulsulfan method. After 6 weeks, chimeric mice (CD45.2 *Irg*1 KO: CD45.1 Wt) were immunized with MOG35-55. At the peak of disease, the mice were sacrificed, the brain/spinal cords were harvested, and single-cell suspensions were prepared via the Percoll density‒gradient method. **B.** CD45.1 and CD45.2 cells were gated and FACS-sorted. The sorted populations were rerun on a BD FACSAria™ III to check the purity of the sorted population. **C.** RNA-seq was performed on CD45.1- and CD45.2-sorted cells from 5 independent animals. Heatmap representing RNA-seq gene expression in the CD45.1 and CD45.2 groups (n=5 animals, padj>0.1 and logFC<0.58). **D.** Volcano plots showing the LogFC versus - log10 p value of CD45.2 in comparison to that of CD45.1. The dots in blue represent upregulated genes, those in gray represent non significantly upregulated genes, and those in red represent upregulated genes. **E.** Normalized expression counts for the specific *Irg1* gene and Irg1-dependent genes (Nrf2 and Hmox1) are shown along with the expression of the Cd45 gene (**F**) as a bar graph (Student’s t test, unpaired) *p < 0.05, **p < 0.01, ***p < 0.001, ns= nonsignificant. **G.** Lollipop plots showing the results of the KEGG analysis, with the length of the lollipops indicating fold enrichment, the color representing -log10(FDR) and the size of the circles representing the number of genes. **H.** Heatmaps showing the logFC values for cell type, cell phenotype markers, macrophage anti- and proinflammatory genes and antigen presentation/processing genes. The color key is shown on the side. Significance is shown on the side of each gene. *p < 0.05, **p < 0.01, ***p < 0.001, ns= nonsignificant.

For functional analysis, we performed Kyoto Encyclopedia of Genes and Genomes (KEGG) pathway analysis. Interestingly, the upregulated genes were enriched in pathways related to T-cell immune functions such as Th17 cell differentiation, TCR signaling, Th1 and Th2 cell differentiation, NK cell-mediated cytotoxicity, hematopoietic cell lineage, cytokine‒cytokine receptor interaction and NFKB signaling (**Fig. 6G**). RNA-seq analysis revealed significant upregulation of the expression of IL17a, RORγt and GM-CSF in *Irg1*-KO cells compared with Wt CD45.1 cells, suggesting that infiltrating mononuclear cells are more pathogenic in nature. Moreover, the higher expression of Class II-associated genes in *Irg1* KO macrophages further suggests that *Irg1*-deficient cells have greater antigen presentation efficacy than Wt cells do. Our Chimera-RNAseq data further support that the expression of anti-inflammatory-associated genes such as Arg1, PPARγ, Msr1, and Mrc1 was significantly lower in CD45.2 KO cells than in CD45.1 wt cells (**Fig. 6H**). These data show that the loss of *Irg1* significantly impacts the expression of key genes involved in a variety of cellular pathways, such as proliferation, differentiation, activation and antigen presentation.

Since *Irg*1 catalyzes the conversion of aconitate into itaconate, we further examined whether supplementation with itaconate could reverse the clinical symptoms of *Irg*1-KO mice. We used dimethyl itaconate (DMI), which has previously been reported to protect against EAE(13). We found that, compared with vehicle-treated Wt EAE mice, DMI-treated Wt EAE mice presented late onset of disease as well as less severe disease (**Supplementary Fig. 8A**). Interestingly, DMI treatment failed to abrogate EAE disease progression in *Irg1* KO mice (**Supplementary Fig. 8A**). We examined the levels of itaconate in DMI-treated animals and found that the levels of itaconate were significantly increased in the blood and other tissues, including the liver, spinal cord and spleen, in the Wt EAE group (**Supplementary Fig. 8B**). In Irg1 KO mice, no itaconate was detected in blood or tissues; however, DMI treatment increased itaconate levels in *Irg*1 KO mice. DMI treatment significantly reduced the number of infiltrating CD4+ T cells expressing IL17a, IFNγ and GM-CSF in the WT EAE mouse group; however, in the Irg1-KO group, the infiltration of CD4+ T cells expressing IL17a was reduced without affecting IFNγ or GMCSF expression (**Supplementary Fig. 8C**). Overall, these sets of data suggest that Irg1 could play a protective role in autoimmune disease without involving itaconate, which needs further study.

## DISCUSSION

The abundance of pathogenic CD4 cells is an important characteristic of both MS and EAE (45). However, the process by which naïve CD4 T cells transform into pathogenic Th17 cells remains unclear. The results of the present study provide evidence that *Irg1*, a mitochondrial enzyme responsible for the production of itaconate, plays a significant role in the pathogenicity of Th17 CD4 cells in EAE. There are few studies reporting the immunoregulatory impact of *Irg1*(8, 46), and our present study demonstrated that *Irg1* is an important regulator of natural defense mechanisms. However, recent studies have shown that the introduction of itaconate can reduce the course of EAE (13, 14). Nevertheless, the extent to which *Irg1* contributes to the control of immune cells in the context of EAE disease remains uncertain.

*Irg1* is a mitochondrial enzyme that is highly upregulated during infection (8), which is consistent with our observation of increased expression at both the mRNA and protein levels in several types of activated immune cells, such as CD11b myeloid cells, monocytes, B cells, and CD4 T cells. Itaconate, a product of *Irg1*, has been shown to provide protection against EAE disease by reducing neuroinflammation(13) and altering the metabolic and epigenetic programming of T cells(14). While the impact of itaconate on the metabolic regulation of Th17 and Treg balance has been documented, the expression of *Irg1* in CD4+ T cells have not been documented. The majority of related studies have focused mostly on the myeloid-centric function of *Irg1* in inflammation. However, our findings indicate that its expression is elevated in CD4 and B cells during EAE compared with CFA. This raises the question of whether *Irg1* has a protective or harmful role in autoimmune diseases such as MS. In the future, we will investigate the cell-specific role of *Irg1* in CNS autoimmunity in CD4+ or B cells. However, our findings indicate that *Irg1* plays a crucial role in regulating macrophages, which are antigen-presenting cells with a hyperinflammatory phenotype. Compared with Wt mice with EAE, *Irg1* knockout mice subjected to EAE induction presented heightened inflammation and demyelination. Alterations in immune cells, including altered phenotypic states, functional states and altered metabolic states, are major causes of EAE progression (45). The metabolic alterations in the macrophages of CD4+ T cells were not the focus of this investigation. However, future studies should be designed to specifically investigate these changes.

Given that the infiltration of macrophages into the central nervous system (CNS) is a key characteristic of multiple sclerosis (MS)(45, 47), our observations indicate a noteworthy increase in the infiltration of circulating monocytes and macrophages into the CNS in *Irg1*-KO mice compared with Wt mice. These findings suggest that the expression of *Irg1* plays a crucial role in the infiltration of immune cells in EAE. In addition, *Irg1*-KO mice exhibited severe disease, and macrophages lacking *Irg1* displayed increased production of proinflammatory cytokines, suggesting that *Irg1* impacts the functional states of immune cells and is involved in EAE pathogenicity. Our results align with those of previous studies, which revealed that macrophages lacking *Irg1* exhibited a proinflammatory macrophage phenotype (8, 44, 48).

Our study revealed a significant mechanism by which *Irg1* suppresses the inflammatory response and the ability of macrophages to present antigens. This is achieved through the activation of the NLRP3–IL-1β cascade, which regulates pathogenic Th17 cells and ultimately controls CNS autoimmunity. However, we have not investigated the role of Irg1-mediated metabolic regulation in this mechanism. The induction of *Irg1* in different cell types during EAE appears to be an innate defensive mechanism aimed at regulating inflammation and the course of the disease. While the levels of itaconate are significantly increased in immune cells and in the spinal cord during EAE, it remains uncertain whether its concentration exceeds the threshold required to effectively regulate inflammation and restore mobility. It has been reported that itaconate can increase the level of IFN-β. Additionally, *Irg1* KO macrophages produce significantly lower levels of IFN-I and its downstream genes related to type I IFN signaling than Wt macrophages do (49). These findings suggest that itaconate could be another contributing factor to the increased severity observed in *Irg1* KO mice with EAE, which has not been thoroughly investigated.

In the present study, we observed that the loss of *Irg1* led to aberrant expression of double-positive (IL17a^+^IFNγ^+^) and triple-positive (IL17A^+^IFNγ^+^GM-CSF^+^) cells, which are known to constitute an antigen-specific CD4 T-cell population and are highly pathogenic in nature(50, 51). Antigen presentation is one of the crucial mechanisms involved in the CD4^+^ T-cell-derived immune response (52); although the pathophysiology of MS is still not entirely understood, it is well known that antigen presentation and the immune response play critical roles in EAE disease progression (45, 47, 53). Our studies *in which Irg1-*KO macrophages were cocultured with MOG-primed CD4 cells clearly demonstrated that compared with Wt macrophages, macrophages lacking *Irg1* expression promoted the development of a pathogenic Th17 phenotype. Th17 cells, a subset of T helper cells that contribute to the immune response against infections and autoimmunity(54), can be categorized as either pathogenic or nonpathogenic on the basis of the type of response they generate and the cytokines with which they interact. Pathogenic Th17 cells, which are mainly mediated by the cytokines IL-1β and IL23, play a role in the development of MS by generating proinflammatory cytokines and recruiting other immune cells to the site of inflammation, whereas nonpathogenic Th17 cells, which are devoid of the involvement of IL-1β and IL23, exhibit a mild inflammatory response. We observed that the splenic macrophages of *Irg1*-KO mice presented increased expression of pathogenic Th17 genes but decreased expression of nonpathogenic Th17 genes, indicating that the expression of *Irg1* may play a regulatory role in controlling the differentiation of Th17 cells. Our research revealed a significant increase in the population of IL-1β-positive cells among both resident and infiltrating macrophages in *Irg1*-KO mice compared with those in WT mice. IL-1β, which is produced by myeloid cells, is a critical cytokine for the development of pathogenic Th17 cells (55, 56). Our findings are consistent with earlier reports that have extensively documented the significant involvement of IL-1β in CNS inflammation, which has been implicated in the progression of neurological disorders such as multiple sclerosis and Alzheimer’s disease (57–59). Immune cells, including macrophages, microglia and astrocytes, produce IL-1β (60), which activates resident immune cells, recruits immune cells from the periphery, and increases the production of additional proinflammatory cytokines in the CNS, thus amplifying the inflammatory reaction.

The production of cleaved IL-1β is regulated by the cleavage of procaspase-1 into active caspase-1, which is mediated by activated NLRP3(61). Compared with that in Wt macrophages, the increased expression of IL1β in *Irg1*-deficient macrophages is consistent with elevated levels of NLRP3 at both the mRNA and protein levels, suggesting that *Irg1* functions in regulating the NLRP3 inflammasome, which is corroborated by earlier findings (42, 62). Antigen presentation and NLRP3 activation are strongly correlated, leading to the regulation of CD4+ T and CD8+ T-cell immune responses (63–65). Our findings indicate that IL-1β alone can induce the development of pathogenic Th17 cells in *Irg1*-KO cells. Additionally, blocking IL-1β limits the differentiation of CD4 cells into pathogenic Th17 cells, suggesting that *Irg1* is involved in the antigen presentation process of *Irg1*-deficient macrophages in the NLRP3-IL-1β-dependent cascade. The aberrant expression of NLRP3 causes severe inflammatory conditions, an increase in the ability of cells to present antigens, dysfunctions in mitochondria, and a failure in resolution processes such as efferocytosis (64, 66). On the basis of our current data, we concluded that *Irg1* regulates Th17-mediated EAE disease progression in a manner that depends on the NLRP3-IL-1β cascade. Collectively, our findings suggest that *Irg*1 expression is a crucial target for regulating the phenotypic and functional activity of both myeloid and lymphoid cells in the progression of EAE, suggesting that targeting *Irg1* could be a promising approach for reducing the severity of MS.

## Materials and methods

### Animals and Induction of EAE

A pair of wild-type (Wt) and *Irg1* KO strains were purchased from Jackson Laboratory and maintained at the Institutional Animal Facility. All animal experiments were approved by the Institutional Animal Care and Use Committee of Henry Ford Health, Detroit, MI. All the mice were housed and bred in a departmental animal facility with controlled humidity and temperature and a 12 h:12 h light/dark cycle with free access to food and water. On day 0, both Wt and *Irg1-*KO mice were immunized via the subcutaneous injection of 200 µl of emulsion containing the MOG35-55 peptide. Pertussis toxin (200 ng/mouse) was injected into the mice on days 0 and 2 postimmunization. Additionally, one separate set of mice (CFA control group) was administered incomplete Freund’s adjuvant (CFA). Disease scoring in the mice was performed in a blinded manner as described previously (15, 67–69) via the traditional EAE scoring method: 0 = no disease, 1 = complete loss of the tail, 2 = partial hind limb paralysis, 3 = complete hind limb paralysis, 4 = complete hind limb and forelimb paralysis, and 5 = death. Detailed Methods and Materials are presented in *<u>SI Appendix, SI Materials and Methods</u>*.

### Statistical analysis

GraphPad version 10 for Windows was used for statistical analysis. All data are expressed as the mean ± standard deviation (SD), and one-way analysis of variance (ANOVA) and t tests were used to determine statistical significance. Statistical significance was defined as a p value of less than 0.05.

## ACKNOWLEDGMENTS

This work is supported in part by research grants from the US National Institutes of Health (AI144004, NS112727), National Multiple Sclerosis Society (US) (RG-2111-38733), and Henry Ford Hospital Internal Support (A10270, A30967) to SG. The funders had no role in the study design, data collection, interpretation, or decision to submit the work for publication.

## AUTHOR CONTRIBUTIONS

MN performed the experiments and wrote the manuscript; MF, FR, KA, MEA performed the experiments and finalized the manuscript; SM, IZ and RR analyzed the data and finalized the manuscript; SG conceived the idea; directed the study; designed the experiments; and finalized the manuscript.

