## Supplementary material for "Immune-responsive gene-1: The mitochondrial Key to Th17 Cell Pathogenicity in CNS Autoimmunity": Supp data

Paste corresponding author: Shailendra Giri

Supp Figure 1

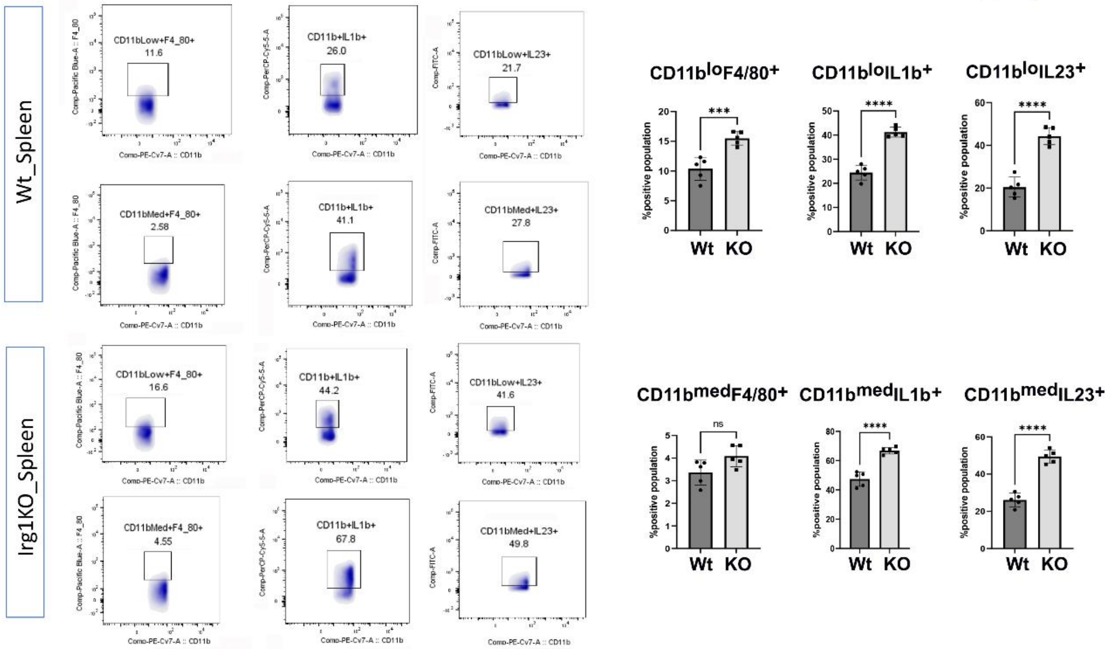

**Supplementary Figure 1. Identification of inflammatory status of Splenic cells in *Irg1* KO mice using flow cytometry.**

Using Classical flow cytometry analysis, the macrophages expressing IL1β and IL-23 in spleen of Wt and KO were characterized. Splenic cells first gated on $CD11b^{low}$ and $CD11b^{med}$ populations, and the total positive populations of F4/80+, IL1β+ and IL23+ were analyzed. Data is presented as mean + SD of 5 animals. **p<0.001, ***p<0.0001, ns = non-significant compared to *Irg1* KO group using student's t test, one-way ANOVA.

**Supp Figure 2**

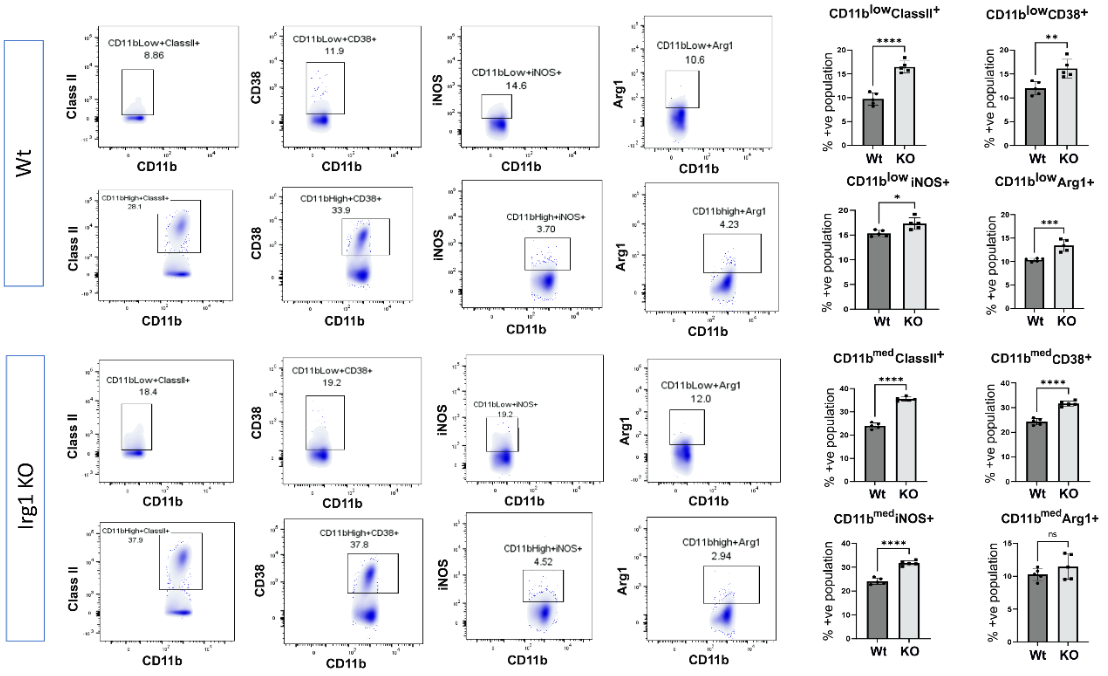

**Supplementary Figure 2. Characterization of polarizations status of splenic cells of *Irg1* KO EAE mice.**

Flow cytometry analysis was done on splenic cells to identify the inflammatory status in Wt and KO EAE mice. Cells were gated on CD11b$^{low}$ and CD11b$^{med}$ and total positive populations of ClassII+, CD38+, iNOS+ and Arg1+ were analyzed. Data is presented as mean + SD of 5 animals. * p<0.01, **p<0.001, ***p<0.0001, ns = non-significant compared to *Irg1* KO group using student's t test, one-way ANOVA.

**Supp Figure 3**

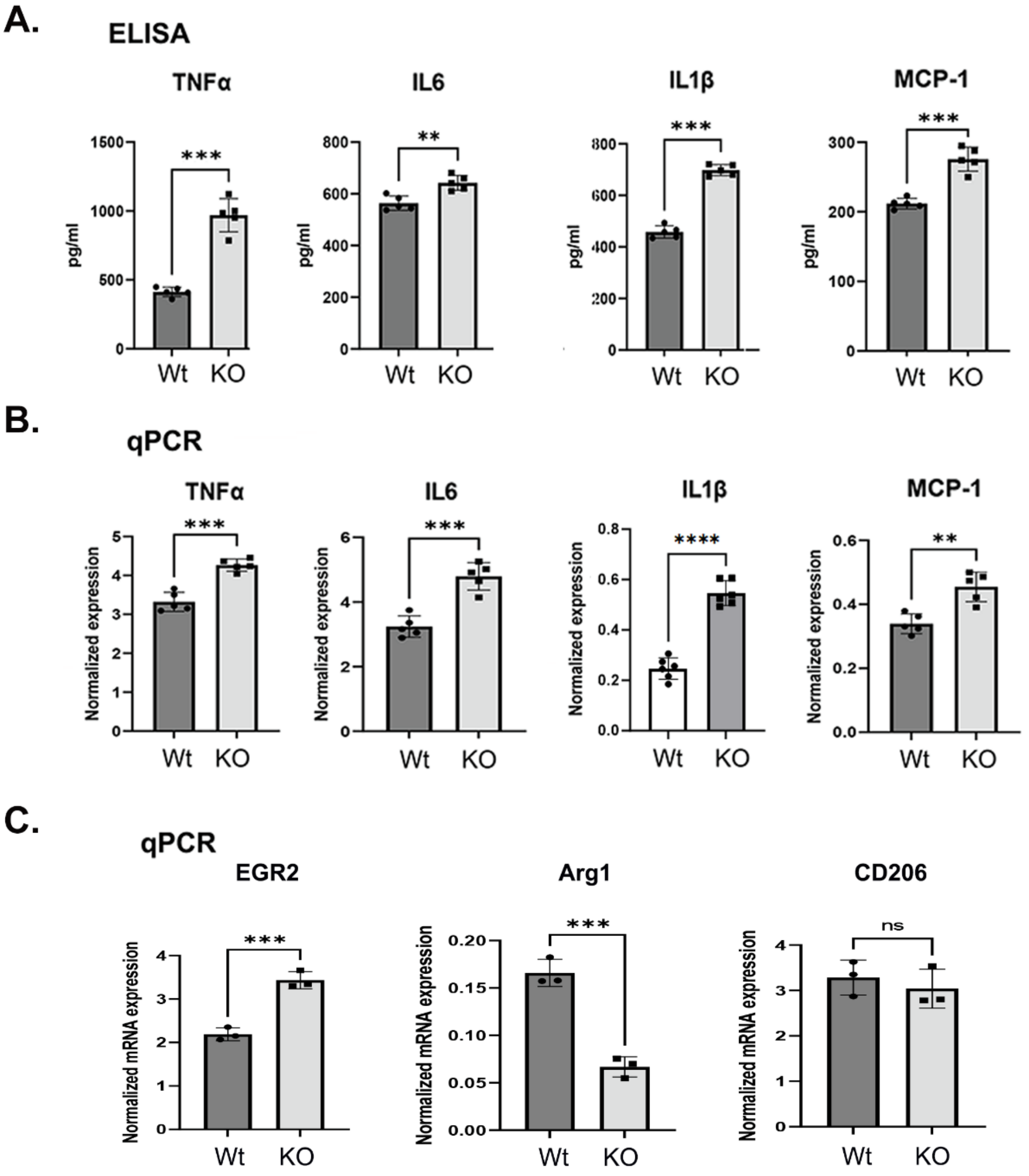

**Supplementary Figure 3. Analysis of M2 genes expression in *Irg1* KO BMDMs.**

The Relative expression of M2 genes (EGR2, Arg1, and CD206) were measured by real time PCR and normalized with the β-actin. Data is presented as mean + SD of 3 animals. **p<0.01, ***p<0.0001, ns=non-significant compared to *Irg1* KO group using student's t test, one-way ANOVA.

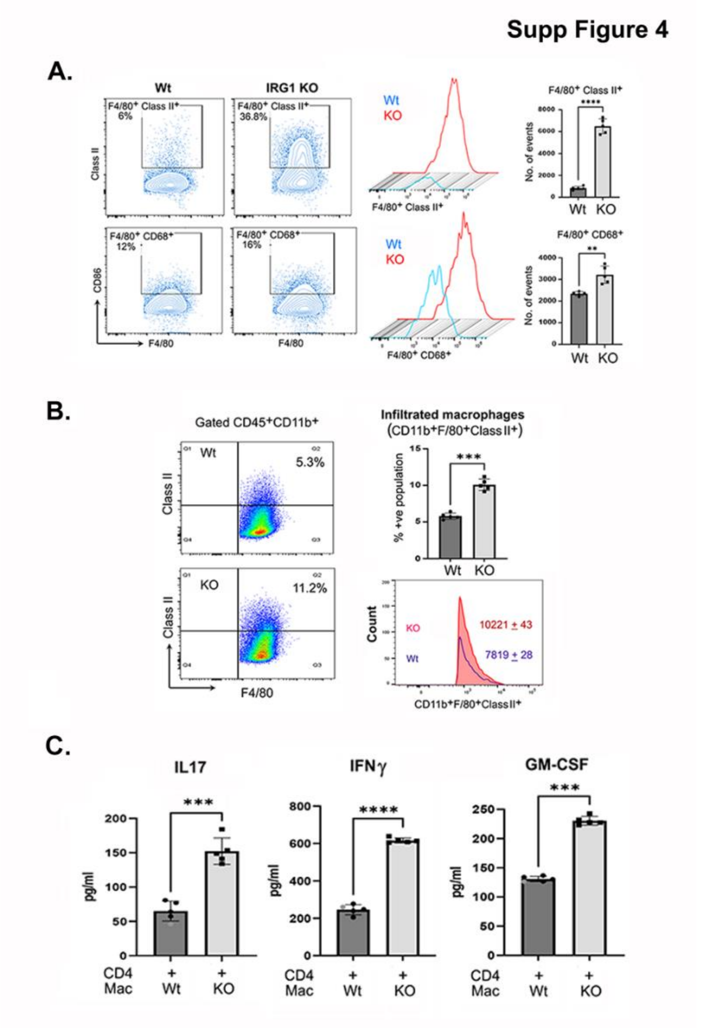

**Supplementary Figure 4. *Irg*1 lacking macrophages promotes antigen presentation capability.**

**A.** Flow cytometry plots (left) of Class II$^{+}$ and CD86$^{+}$ populations in BMDM of Wt and KO upon IFNγ stimulation for 24h and quantifications of Class II$^{+}$ and CD86$^{+}$ cells with histogram overlay (right) of Class II$^{+}$ and CD86$^{+}$ cells between Wt and *Irg1* KO (n=5). **B.** Flow cytometry plot gated (left) on CD45$^{hi+}$CD11b$^{+}$F4/80$^{+}$ (infiltrated macrophages) showing Class II+ populations in Wt and *Irg1* KO and quantifications (right) of Class II$^{+}$ cells with histogram overlay of Class II$^{+}$ cells between Wt and *Irg1* KO (N=5). **C.** Bar graph represents data from ELISA assay demonstrating levels of pro-inflammatory cytokines released by Wt CD4 from co-culture study using Wt CD4 cells with Wt and *Irg1* KO macrophages (N=5) as shown in figure 5H. **p<0.01, *** or ****p<0.0001, ns=non-significant compared to *Irg1* KO group using student's t test, one-way ANOVA.

**Supp Figure 5**

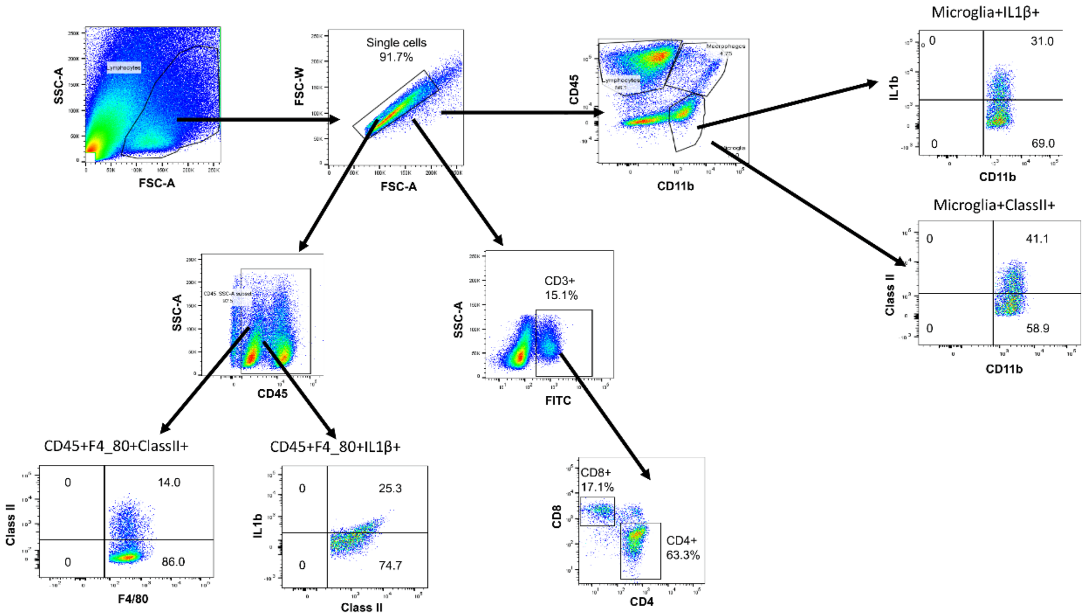

**Supplementary Figure 5: Gating strategy for characterization of different types of Immune cells in the CNS.**

Flow cytometry was used to identify CNS (brain and spinal cord) immune cells in single cells isolated from immunized Wt and *Irg*1 KO mice. $CD45^{low}CD11b^{+}$ populations were gated on single cell to identify microglial cells, and then $IL1\beta^{+}$ and $ClassII^{+}$ populations were analyzed. For characterization of CNS infiltrating cells, $CD45^{hi}CD11b^{+}$ populations were identified. To check the expression of IL1β and ClassII on infiltrated macrophages, $CD45^{+}F4/80^{+}$ populations were gated. The total CD4 population was identified using $CD45^{+}CD3^{+}CD4^{+}$ gating strategy.

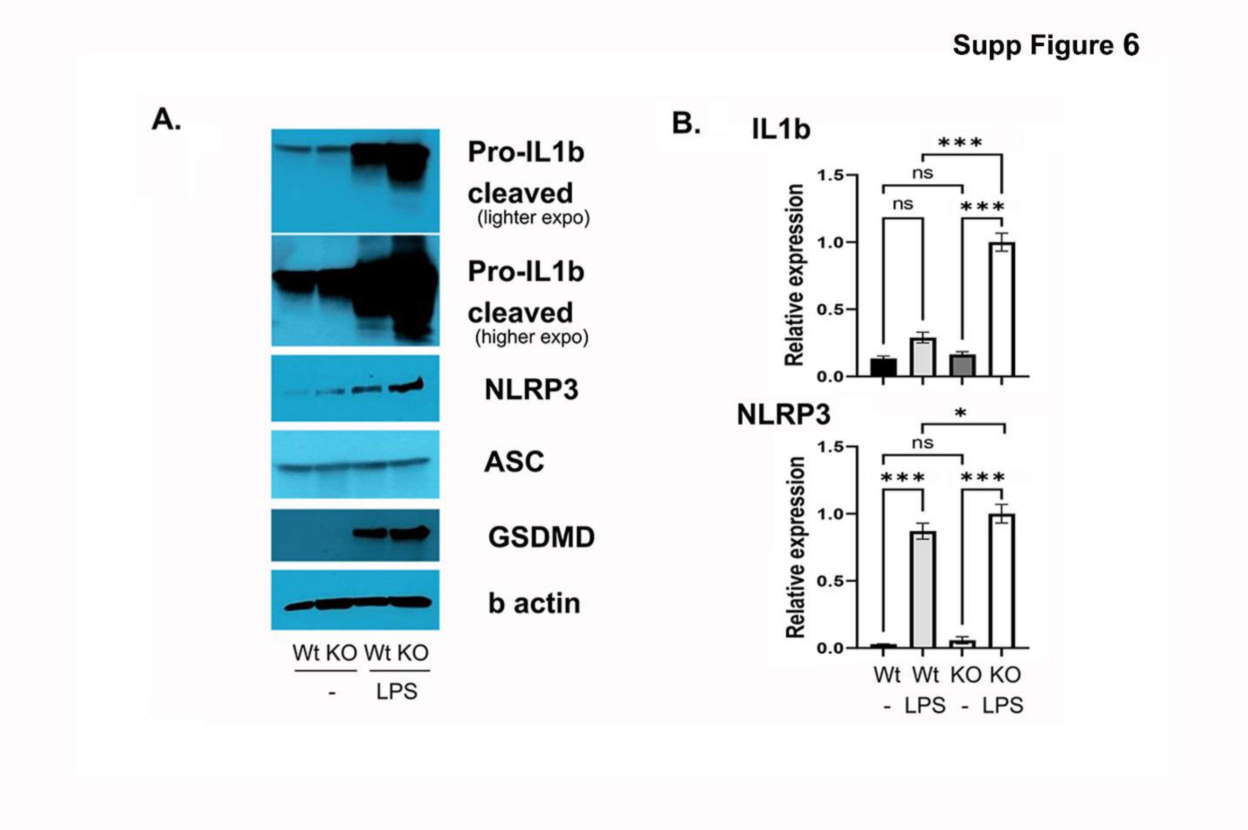

**Supplementary Figure 6: *Irg*1 lacking macrophage produced higher levels of IL1β via NLRP3**. **A.** Immunoblotting of Wt and *Irg1* KO macrophage stimulated with LPS for 4hr showing NLRP3, pro-IL1β, cleaved IL1β, ASC and GSDMD normalized with β-actin. **B.** Real-time quantitative PCR analysis of the fold change of NLRP3 and IL1β mRNA expression upon stimulation with LPS for 4hrs (N=3). *p < 0.05, ***p < 0.001, ns= non-significant.

Supp Figure 7

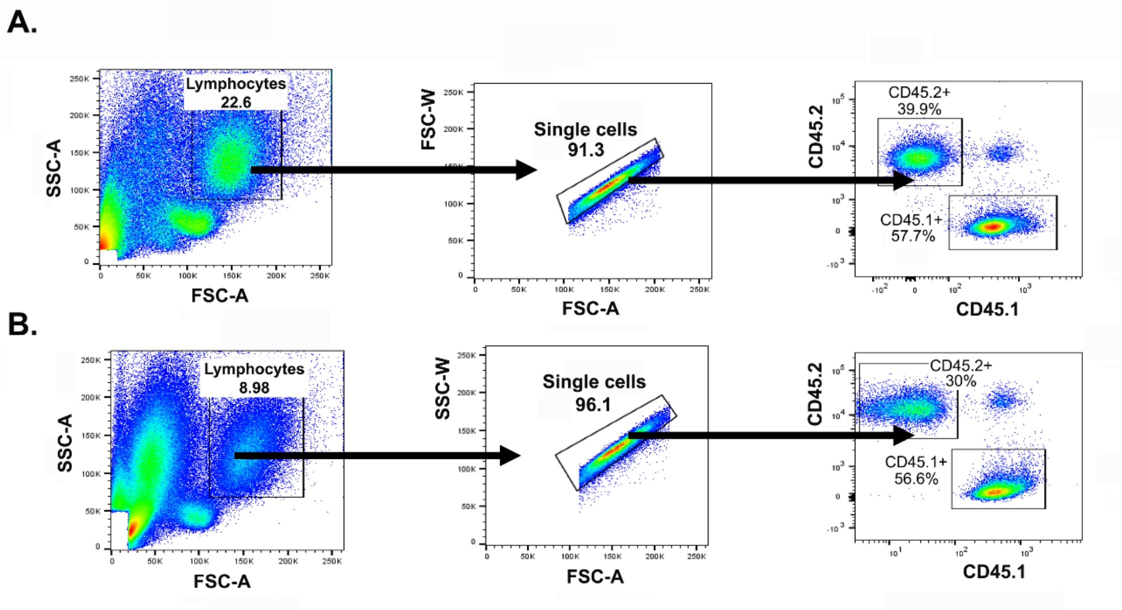

**Supplementary Figure 7: Validation of BM Chimeras**. Prior to EAE Immunization, we harvested spleen (**A**) and blood (**B**) from Chimera mice and examined the populations of CD45.1 and CD45.2 to confirm the development of Chimeric mice model using 50:50 BM of Wt and *Irg1* KO. The distinct populations of CD45.1 and CD45.2 populations represented by classic Flow cytometry (N=5).

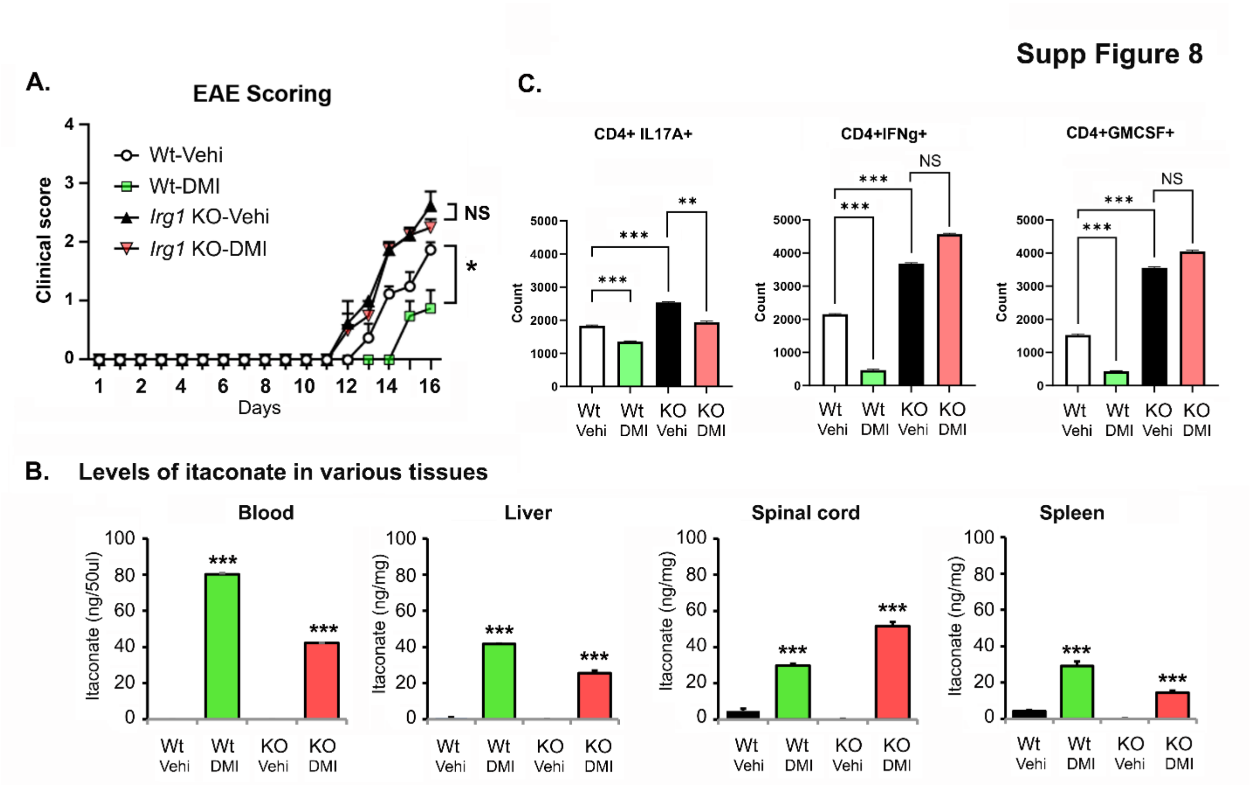

**Supplementary Figure 8: Dimethyl itaconate (DMI) did not provide protection in *Irg*1 KO with EAE. (A)** Graph showing EAE scoring of four different mouse groups; Wt-EAE treated with vehicle (corn oil); Wt-EAE treated with DMI, *Irg*1 KO-EAE treated with vehicle and *Irg*1 KO-EAE treated with DMI. DMI at the dose of 400mg/kg body weight was given daily in corn oil via ip. EAE scores were recorded daily till end of the study. Data are presented as mean ± SEM (N=4). *p < 0.05, NS= non-significant. **(B)** Levels of itaconate were examined in blood, liver, spinal cord and spleen in Wt and *Irg1* KO EAE groups treated with Vehicle or DMI. Data are presented as mean ± SEM (N=3). **(C)** BILs were profiled with gated CD45+CD4+ cells expressing IFN□, IL17a and Gmcsf. Data are presented as mean ± SEM (N=3). NS not significant; ***p < 0.001.

**Supplementary Table 1**

| Antibodies | Catalogue No. | Manufacturer's Name |
|---|---|---|
| CD16/32 | Biolegend | 101302 |
| AF647-Puromycin | Biolegend | 381508 |
| BV421-CD45 | Biolgend | 103134 |
| BV510-CD11b | Biolegend | 101263 |
| FITC-CD11b | Biolegend | 101206 |
| PE-Cy7-CD11b | Biolegend | 101216 |
| BV510-Ly6G | Biolegend | 127633 |
| PE/Dazzle594-Ly6G | Biolegend | 127648 |
| AF700-Ly6C | Biolegend | 128024 |
| Percp/Cy5.5-CD45 | Invitrogen | 103132 |
| PE-Cy5.5-CD45 | Biolegend | 35-0451-82 |
| BV421-F4_80 | Biolegend | 123132 |
| APC/Cyanine7-CD11c | Biolegend | 117324 |
| BV510-ClassII | Biolegend | 107636 |
| BV421-CD38 | Biolegend | 102732 |
| FITC-iNOS | BD Biosciences | 610330 |
| PE-Arg1 | Biolegend | 165804 |
| Percp-eFluor 710-IL1β | eBiosciences | 46-7114-82 |
| AF488-IL23 | Invitrogen | 53-7023-82 |
| PE-GMCSF | Biolegend | 505406 |
| Texas Red-IL17a | Biolegend | 506938 |
| PE-Cy7-Tbet | Biolegend | 644824 |
| BV421-RORγt | BD Biosciences | 562894 |

**Supplementary Table 2**

| Oligonucleotides | Source | Forward & Reversed Sequence |
|---|---|---|
| m*Irg*1 | Realtimeprimers.com | Fw: GGT ATC ATT CGG AGG AGC AAG AG<br>Rv: ACA GTG CTG GAG GTG TTG GAA C |
| r*Irg1 (rat)* | Bio-Rad | Cat: 10041596 |
| mILβ | Integrated DNA technology (ITD) | Fw: TGG AAA AGC GGT TTG TCT TC<br>Rv: TAC CAG TTG GGG AAC TCT GC |
| mTNFα | Integrated DNA technology (ITD) | Fw: GGT GCC TAT GTC TCA GCC TCT T<br>Rv: GCC ATA GAA CTG ATG AGA GGG AG |
| mIL6 | Integrated DNA technology (ITD) | Fw: GAG GAT ACC ACT CCC AAC AGA CC<br>Rv: AAG TGC ATC ATC GTT GTT CAT ACA |
| mNLRP3 | Integrated DNA technology (ITD) | Fw: TCA CAA CTC GCC CAA GGA GGA A<br>Rv: AAG AGA CCA CGG CAG AAG CTA G |
| mMCP-1 | Integrated DNA technology (ITD) | Fw: GAG AGC TAC AAG AGG ATC ACC A<br>Rv: GTA TGT CTG GAC CCA TTC CTT C |
| mRORγt | Integrated DNA technology (ITD) | Fw: CCG CTG AGA GGG CTT CAC<br>Rv: TGC AGG AGT AGG CCA CAT TAC A |
| mIL17a promoter | Integrated DNA technology (ITD) | Fw: TAATTCCCCCACAAAGCAAC<br>Rv: TGAGGTCAGCACAGAACCAC |
| Mouse ribosomal protein L27 | Realtimeprimers.com | Cat: VMPS-5561 |
| Rat ribosomal protein L27 | Realtimeprimers.com | Cat: VRPS-5294 |

### SI Materials and Methods

#### Primary brain rat glial culture

The timed pregnant rat was purchased from Charles River Laboratory. The brains of 2–3-day-old female and male pups were used to prepare mixed glial cell cultures. We followed our previously published protocol to prepare primary mixed glial cell cultures [1]. After 10 days of culture, the cells were treated with LPS (0.5 μg/ml) + IFNγ (20 ng/ml) to induce inflammatory conditions in our study. After 8 h of treatment, the cells were processed for RNA isolation and RNA-seq as described below.

#### Bone marrow chimeras and FACS sorting

For the generation of bone marrow chimeras, host (CD45.1) animals were treated with three doses of busulfan (20 mg/ml), and on the fourth day, the host received an i.v. injection of 10 million BM cells [50:50 mixture of CD45.1 (Wt) and CD45.2 (*Irg1* KO)]. Using flow cytometry, we sorted CD45.1+ and CD45.2+ cells from BILs isolated via a Percoll gradient from the brain and spinal cord after EAE induction. BILs were stained with primary antibodies against mouse PerCP-Cy5.5-CD45.1 and BV421-CD45.2 (BioLegend) and sorted on a BD FACSAria™ III. Furthermore, the sorted populations were re-run to check the purity of the sorted cells. Next, we used an automated cell counter (Bio-Rad) to determine the viability of CD45+1+ and CD45.2 cells via trypan blue and found that >98% of the cells were viable. Then, RNA was harvested from the sorted cells and processed for RNA sequencing (**Supplementary Fig 7**).

#### RNA-seq

We extracted total RNA from rat mixed glial cells stimulated with LI for 8 h in serum-free DMEM; three replicates were used with a Qiagen RNeasy Kit. Similarly, for the CD45 samples, we isolated total RNA from half a million CD45.1- and CD45.2-sorted cells with a Qiagen RNeasy Mini Kit. The samples with a high RIN of more than 7.28 were selected for library preparation for RNA-seq. PolyA-enriched RNA libraries were prepared and barcoded for multiplexing according to Illumina recommendations. Paired-end sequencing (150 bp) was performed with an Illumina NovaSeq 6000 via NOVOGENE INC. We performed QC with FastQC and aligned 150 bp paired-end reads to GRCm38-mm10, with approximately 80% uniquely mapped reads in mixed glial samples and 89%-94% in CD45 samples via the STAR aligner. The STAR aligner was subsequently utilized for count calling for each sample. For differential expression analysis, we employed the limma-Voom package in R, with a padj cutoff of 0.0005 and a log-fold change (logFC) threshold of 1.5 in mixed glial samples, and in CD45 samples, a padj cutoff of 0.1 and logFC of 0.58 (FC=1.5) were set. The threshold for filtering out genes with low expression was set to 0.1 counts per million (CPM) in mixed glial samples and 0.5 CPM in at least 80% of the CD45 samples. For count normalization, we used the trimmed mean of M values (TMM) method. Finally, R was used to generate the plots for visualization.

**Single-cell RNA sequencing**

Single-cell RNA capture, with a total of 0.2 million cells, was performed via the PIPseq T20 3' single-cell capture and lysis kit. Three sc-RNA libraries from each group were prepared with a fluent PIPseq T20 3' single-cell RNA kit according to the manufacturer's instructions. The libraries were sequenced on a partial lane of an Illumina NovaSeq X–Plus sequencer. The 150 bp paired-end reads were then aligned to the mm10 mouse genome with Pipseeker v3.0 software. Following barcode matching, count matrices were generated, which were piped with Seurat V3 for further analysis. Individual seurat objects were created, with QC parameter filters of a minimum of 200–600 features and < 5% mitochondrial genes, which were merged later according to the groups. Next, approximately 16K genes across ~8000 single cells were analyzed for subsequent analysis, including normalization, scaling, and dimensionality reduction using Principal Component Analysis (PCA) and Uniform Manifold Approximation and Projection (UMAP), as implemented in the Seurat pipeline. We utilize canonical correlation analysis (CCA) methods for identifying anchors to integrate the two groups of scRNA-seq datasets through the application of the FindIntegrationAnchors function. The preprocessed data were subjected to clustering analysis via Seurat's FindNeighbors and FindClusters functions, with the first 30 significant PCs and a resolution of 0.25. The marker genes for each cluster were identified via the FindAllMarkers function. The cell types were annotated on the basis of the expression patterns of these markers, referring to known available cell type information. A heatmap was generated via the DoHeatmap function.

**Immunohistochemistry**

The mice were anesthetized with isoflurane and transcardially perfused with 0.9% saline followed by 4% paraformaldehyde in 0.1 M phosphate buffer. The brains were removed, postfixed overnight in paraformaldehyde and cryoprotected in 30% sucrose at 4°C until they sank. Coronal sections (25 µm) were cut on a cryotome (Thermo CS3472M0402; Warrington UK), and tissue was collected for histological assessment. Diaminobenzidine (DAB) immunostaining was performed by blocking the sections with 1% BSA in PBS for 1 h at room temperature, followed by incubation with a polyclonal anti-*Irg1* antibody (1:250, CST 19857, Cell Signaling Technology) overnight at 4°C. The sections were washed three times with 0.1% PBST. The sections were washed and incubated with a biotinylated secondary antibody for 30 min at room temperature. The sections were subsequently developed via VECTASTAIN Elite ABC system according to the manufacturer's instructions (Vector Laboratories, Burlingame, CA). The sections were washed three times, mounted with mounting medium and covered with a glass cover slip. The images were scanned with Aperio eSlide Manager (Leica Biosystems). Quantification was performed by using ImageJ software (Version 1.49; NIH, USA). The data are presented as the means ± SEMs.

**Isolation and characterization of Brain Infiltrating Cells (BILs)**

A Percoll density gradient was used to isolate BILs from CNS tissues (brain and spinal cord), as previously described[1-4] with minor modifications. Briefly, single-cell suspensions were prepared from CNS tissues via mechanical digestion via 18G and 22G needles, and enzymatic dissociation was performed via Accumax (Innovative Cell Tech). Then, the brain homogenates were passed through a 100 µM strainer to obtain single-cell suspensions. A 30% Percoll density gradient was used to remove myelin, and mixed glial populations were collected from pellets. The cells were washed with PBS and used for further experimental analysis. To characterize the phenotypic and functional states of immune cells, a flow cytometry assay was performed. The collected cells were resuspended in 500 µL of FACS stain buffer. The cells were washed three times and resuspended in 100 µl of FACS stain buffer. To block nonspecific binding, CD16/32 was added to the samples, which were subsequently incubated on ice for 5 min in the dark, followed by the addition of a number of surface antibodies, as indicated for myeloid and lymphoid cells, for 30 min at 4°C in the dark for flow cytometry. After incubation, the cells were washed with FACS stain buffer, fixed and permeabilized via the Fixation/Permeabilization Buffer Set (Proteintech), following the manufacturer's instructions. After this, intracellular staining of the antibodies (**supplementary table 1**) directed against the functional states of myeloid and lymphoid cells was performed in permeabilization buffer for 1 h at room temperature. After incubation, the cells were washed 3 times with FACS buffer, run on a Thermo Fisher Attune cytometer and analyzed via FlowJo software. The flow cytometry gating strategy is described in the supplementary material (**Supplementary Fig. 5**).

#### RNA isolation and quantitative PCR analysis

TRIzol reagent was used to isolate RNA from the spinal cord, spleen and bone marrow-derived macrophages (BMDMs) according to the manufacturer's instructions. The RNA concentration and purity were determined via a Nanodrop spectrophotometer. The RNA was reverse transcribed via the iScript cDNA Synthesis Kit (Bio-Rad) according to the manufacturer's instructions. Using specific primers, the mRNA expression of various genes associated with functional changes associated with immune cells and primers directed against the *Irg1* gene were quantified. The primer list is shown in the table (**supplementary table 2**). A CFX Real-Time PCR Cycler (Bio-Rad) was used to perform real-time qPCR. Using L27 as an internal standard, relative mRNA expression was calculated via CFX Maestro software (Bio-Rad), and the relative amount of mRNA is presented as the fold change over the control.

#### Immunoblotting analysis

The spinal cord, splenic, BMDM and different myeloid and lymphoid cells were lysed via protein lysis buffer containing a protease inhibitor cocktail. After 30 min of incubation, the samples were centrifuged at 10,000 × g for 20 min at 4°C. Protein estimation was performed via the Bradford assay. Thirty micrograms of protein were loaded on a 4–20% gradient Bio-Rad precast gel. The proteins were subsequently transferred to nitrocellulose membranes. After transfer, the membrane was blocked with 5% skim milk in TBST and incubated with the indicated primary antibodies overnight at 4°C. The next day, the membranes were rinsed three times with TBST and incubated with the corresponding secondary antibody at room temperature for 2 h. After washing with TBST, protein expression was visualized via the addition of enhanced chemiluminescence (ECL) reagent (Thermo), and densiometric analysis was performed via ImageJ software.

#### Adoptive transfer (Adt) and Recall experiments

The AdT experiment was performed as described previously [2]. Briefly, the spleen and LNs were isolated, and single cells were prepared, followed by RBC lysis. Furthermore, single-cell suspensions were restimulated with 20 µg/ml $MOG_{35-55}$ peptide in the presence of anti-IFNγ (10 μg/ml, BioXcell) and IL12p70 (20 ng/ml) for 72 hrs. After incubation, the cells were harvested and washed, and 10 million cells were i.p. injected into B6 Wt mice. On day 0 and day 2, each mouse was administered pertussis toxin. EAE scoring was performed daily through the traditional scoring method. For the adoptive transfer experiment in Rag1 mice, CD4 T cells were isolated from both Wt and *Irg1*-KO mice and restimulated as described above. Then, 10 million CD4+ T cells were injected i.p. into Rag1 mice. For the recall response, spleen and LN cells were isolated and cultured with $MOG_{35-55}$ peptide (20 ng/ml), and the supernatants were collected at 48 hr, 72 hr and 96 hr. The levels of both proinflammatory cytokines and anti-inflammatory cytokines produced were measured via ELISA (BioLegend).

**Analysis of the immune response in *Irg1*-KO mice**

To analyze the inflammatory response, single cells were prepared from the spleen/LNs, and BILs were harvested from CNS tissues. Splenic/LN cells and BILs were restimulated with $MOG_{35-55}$ for 72 hr, and the supernatants were collected for estimation of cytokine production via ELISA. The cells were stimulated with PMA and ionomycin for 4 h. Then, the cells were surface stained with FITC-conjugated anti-CD4 antibodies and intracellularly stained with anti-IL17a, anti-IFNγ, GM-CSF, T-bet, RORγt and IL4 antibodies.

**Generation of bone marrow-derived macrophages (BMDMs)**

Bone marrow-derived macrophages were prepared as described previously[2]. Bone marrow cells were flushed from the femora and tibias of Wt and *Irg1*-KO mice. After centrifugation, the cells were cultured with complete RPMI-1640 containing 10% L-929 media for seven days and then fed complete media. After seven days, the cells were trypsinized, and 1–2 million cells from each group were seeded into a new plate as indicated.

**Chromatin immunoprecipitation (ChIP) assay**

A chromatin immunoprecipitation (ChIP) assay was performed via a Go-Chip-Grade Kit (BioLegend) according to the manufacturer's instructions, with slight modifications. Briefly, 1% formaldehyde was directly added to the cells to cross-link the proteins with the DNA, after which 10% glycine was added to neutralize the reaction. After this, the cell pellets were collected via centrifugation at 2000 rpm for 6 min. The pellets were dissolved in sonication lysis buffer and then broken into 200-bp pieces via Bioraptor plus (Diagenode). The collected fragments were incubated for an hour at 4°C in a shaker with protein A/G magnetic beads (Selleckchem). The mixture was subsequently centrifuged, and the pellets were incubated with either anti-mouse RORγt (Proteintech) or anti-mouse IgG (Proteintech) antibodies at 4°C overnight. After this, protein A/G magnetic beads were added to the antibody–pellet complex for 1 hr at 4°C with shaking. After incubation, the antibody–pellet complex was magnetically purified via a magnetic separator. Then, 30 µL of eluent was added to the antibody–DNA complexes for 30 min. A total of 20 µl of 5 M NaCl was added to the eluted samples, which were subsequently incubated at 65°C overnight to decrosslink the samples. The DNA was then purified via a Biolegend kit. The DNA extracted from the antibody-immunoprecipitated chromatin fragments was subjected to qPCR to amplify the fragment region of the binding sites on the IL17a promoter. The 5′-TAATTCCCCCACAAAGCAAC-3′ (forward) and 5′-TGAGGTCAGCACAGAACCAC-3′ (backward) primers were used to target the IL17a promoter sequences.
